# Shared and specific functions of Arfs 1–5 at the Golgi revealed by systematic knockouts

**DOI:** 10.1101/2021.04.19.440443

**Authors:** Mirjam Pennauer, Katarzyna Buczak, Cristina Prescianotto-Baschong, Martin Spiess

**Affiliations:** Biozentrum, University of Basel, Klingelbergstrasse 70, CH-4056 Basel, Switzerland

**Keywords:** Arf GTPases, Golgi, secretory pathway, COPI, AP1, GGA

## Abstract

The ADP-ribosylation factors (Arfs) are small GTPases regulating membrane traffic in the secretory pathway. They are closely related and appear to have overlapping functions, regulators, and effectors. The functional specificity of individual Arfs and the extent of redundancy *in vivo* are still largely unknown. We addressed these questions by CRISPR/Cas9-mediated genomic deletion of the human class I (Arfs 1 and 3) and class II (Arfs 4 and 5) Arfs, either individually or in combination. Cells lacking individual Arfs or certain combinations were viable with only a slight growth defect when lacking Arf1 or Arf4. However, Arf1 and 4, and Arf4 and 5 could not be deleted simultaneously. Hence, class I Arfs are not essential and Arf4 alone was found to be sufficient for cell viability. Remarkably, two single knockouts produced specific and distinct phenotypes. Upon deletion of Arf1, the Golgi complex was enlarged and recruitment of vesicle coats decreased, confirming a major role of Arf1 in coat formation at the Golgi. Cell lines deleted for Arf4 exhibited secretion of ER resident proteins, indicating a specific defect in coatomer-dependent ER protein retrieval by the KDEL receptors. The knockout cell lines will be a useful tool to study other Arf-dependent processes.

## Introduction

The secretory pathway is a major route of membrane traffic in the cell, transporting soluble and membrane proteins from their site of synthesis, which is the rough endoplasmic reticulum (ER), to their final destinations. On the way, cargo proteins pass through successive compartments, where they acquire modifications and undergo multiple rounds of sorting and packaging into transport carriers. This anterograde traffic is counterbalanced by retrograde transport of membranes and proteins to maintain organelle identity and homeostasis, and retain specific proteins in defined compartments. Key players in these processes are the small GTPases of the Arf (ADP-ribosylation factor) family.

The Arf family comprises 30 members: the six “true” Arfs, 21 Arls (Arf-like proteins), 2 Sars and Trim23 (Li et al 2004, Kahn et al 2006). The Arfs are closely related, while the other members are more divergent in sequence and cellular functions (reviewed in Gillingham and Munro 2007, Donaldson and Jackson 2011, Sztul et al. 2019). The five human Arfs (humans lack Arf2) are assigned to 3 classes based on sequence homology: Arf1 and 3 belong to class I, Arf4 and 5 to class II, and Arf6 is the only member of class III. Class I and II Arfs mainly localize to the Golgi, but also to endosomes and/or the ER-Golgi intermediate compartment (ERGIC), whereas Arf6 is found in the cell periphery. Arfs are ubiquitously expressed, but vary in their abundance (Cavenagh et al. 1996, Itzhak et al. 2016). In the widely used HeLa cells, Arf1 is the most abundant Arf, followed by Arf4 (∼1/3), Arf5 and Arf6 (∼1/10), and Arf3 (∼1/100) (Itzhak et al. 2016).

Arfs are N-myristoylated, which allows them to loosely associate with membranes already in the GDP-bound state. Binding of a guanine nucleotide exchange factor (GEF) and subsequent activation via GDP–GTP exchange lead to displacement of the N-terminal amphipathic helix from the hydrophobic binding pocket, resulting in tight membrane association (Antonny et al. 1997, Renault et al. 2003). Concomitant conformational changes enable binding of effectors.

The interplay of Arfs and their various effectors contribute to diverse cellular processes throughout the cell (reviewed in Donaldson and Jackson 2011, Jackson and Bouvet 2014, Sztul et al. 2019). The most prominent function is the contribution of the Golgi-localized Arfs (Arf1–5) to transport carrier formation in intracellular traffic, especially in the secretory pathway. Two major aspects are linked to Arf activity in this context: the modification of membrane lipids (reviewed in de Matteis and Godi 2004, Donaldson and Jackson 2011) and the recruitment of coat components. The best-characterized coat complexes are the coatomer complex I (COPI) at the ERGIC and the Golgi mainly for retrograde transport back to the ER, and the adaptor protein complex 1 (AP1) and the Golgi-localized, γ-ear-containing, Arf-binding proteins (GGAs) at the Golgi and on endosomes for transport from the trans-Golgi network (TGN) to endosomes and back.

The activity of Arfs is tightly regulated spatially and temporally by their GEFs and GTPase-activating proteins (GAPs). All 15 known GEFs share a common Sec7 domain to catalyze nucleotide exchange, but in addition possess diverse domains regulating their own membrane association and activity (reviewed in Nawrotek et al. 2016, Sztul et al. 2019). Also, the 28 ArfGAPs share a common GAP domain and are increasingly perceived to be more than simple terminators of Arf activity, but rather effectors themselves (reviewed in Donaldson et al. 2011, Sztul et al. 2019).

Originally discovered as a factor required for cholera toxin-mediated stimulation of adenylate cyclase by ADP-ribosylation of the stimulatory heterotrimeric G protein Gs (Kahn and Gilman 1984, 1986), the role of Arfs in intracellular traffic by recruiting coat proteins was uncovered a few years later (Stearns et al. 1990, Serafini et al. 1991, Stamnes and Rothman 1993, Palmer et al. 1993, Traub et al. 1993). Early approaches to identify Arf functions were based on the manipulation by dominant-negative and dominant-active mutants (Teal et al. 1993, Zhang et al. 1994, Dascher et al. 1994) and by the fungal macrolide Brefeldin A (BFA). However, these approaches lacked specificity for individual Arfs, since Arf mutants and BFA sequester shared GEFs or GAPs and hence influence the activity of various Arfs simultaneously. Direct interactions between Arfs and coat components were analyzed in the presence of GTPγS by *in vitro* and *in vivo* experiments, which suggested that both class I and II Arfs can recruit COPI, AP1 and GGAs to Golgi membranes (Liang and Kornfeld 1997, Boman et al. 2000, Austin et al. 2002, Takatsu et al. 2002).

Volpicelli-Daley and colleagues (2005) were the first to systematically dissect the role of individual Arfs in the secretory and endocytic pathway by siRNA-mediated knockdown. They proposed that no single Arf is required for any transport step, since only pairwise knockdowns resulted in specific phenotypes, hence suggesting cooperative action of Arfs and some isoform specificity at certain steps in intracellular transport (Volpicelli-Daley et al. 2005).

Later studies provided a glimpse into the complex interactome surrounding the Arfs involving GEFs, GAPs, effectors, and other GTPases, and highlighted that Arfs do not act in isolation, but in complex networks (reviewed in Donaldson and Jackson 2011, Mizuno-Yamasaki et al. 2012, Baschieri et al. 2012, Thomas et al. 2020). However, fundamental questions remained unanswered, such as the major contributions of individual Arfs, their specificities and redundancies and their regulation and coordination (Sztul et al. 2019).

Here, we revisited basic questions concerning the functions of Arf 1–5 in the secretory pathway using CRISPR/Cas9 genome editing, generating Arf knockout cells by genomic deletion. We found that cells lacking any single Arf are viable, as well as cells deleted for certain double− or triple-combinations. In fact, Arf4 is able to sustain all essential functions in the absence of all other class I and class II Arfs. Yet, we observed distinct phenotypes already in single-knockout cell lines: deletion of Arf1 caused an increased Golgi volume, altered Golgi morphology, and reduced recruitment of vesicle coats to the Golgi, while the knockout of Arf4 produced a specific defect in retrieval of ER resident proteins.

## Results

### Generation of Arf knockout HeLaα cell lines

To characterize specific and redundant functions of Arf1–5 in the secretory pathway, we aimed to delete single or multiple Arf proteins by a CRISPR/Cas9-mediated knockout. Two guide RNAs were designed to delete a genomic region of the respective Arf genes including the start codon and the part of exon 1 encoding the N-terminal myristoylated amphipathic helix.

Initially, we knocked out single Arfs and succeeded to obtain all four knockout cell lines (Arf1ko, Arf3ko, Arf4ko, Arf5ko), indicating that no single Arf is essential for viability and cell growth (Table 1). Based on these single-knockout cell lines, we generated cell lines for double- and triple-knockout combinations. The chronological order of knockouts is indicated in the name of the cell lines. In Arf3+1ko, for example, Arf3 was deleted first, followed by Arf1. Cell lines for four out of six double-knockout combinations (Arf1+5ko, Arf3+1ko, Arf3+4ko, Arf3+5ko) and one out of four triple-knockout combinations (Arf3+1+5ko) were successfully generated, while we were repeatedly unable to obtain Arf1+4ko and Arf5+4ko cell lines (Table 1).

**Table 1:** Arf knockout cell lines. This overview lists all attempted Arf knockout cell lines and indicates successful (highlighted in green) or failed generation (highlighted in red). The order of isoforms in double- and triple-knockouts indicates the temporal succession of Arf deletion.

| Knockout | single | double | triple |
| --- | --- | --- | --- |
| Deleted<br>Arf(s) | 1 | 1+4 | 3+1+5 |
|  | 3 | 1+5 |  |
|  | 4 | 3+1 |  |
|  | 5 | 3+4 |  |
|  |  | 3+5 |  |
|  |  | 5+4 |  |

For all work presented here, one representative clonal cell line was carefully chosen for each knockout after initial verification. This included validation of the deletions in all alleles by genomic PCR (Figure 1A) and confirmation of the knockout on the protein level by immunoblot analysis (Figure 1B). In general, the loss of Arf proteins did not strongly affect the protein level of the remaining Arfs, with the exception of Arf4, which was reproducibly upregulated in cells lacking Arf1 by an average of 4.1-fold (±0.9; n=3).

**Figure 1:**
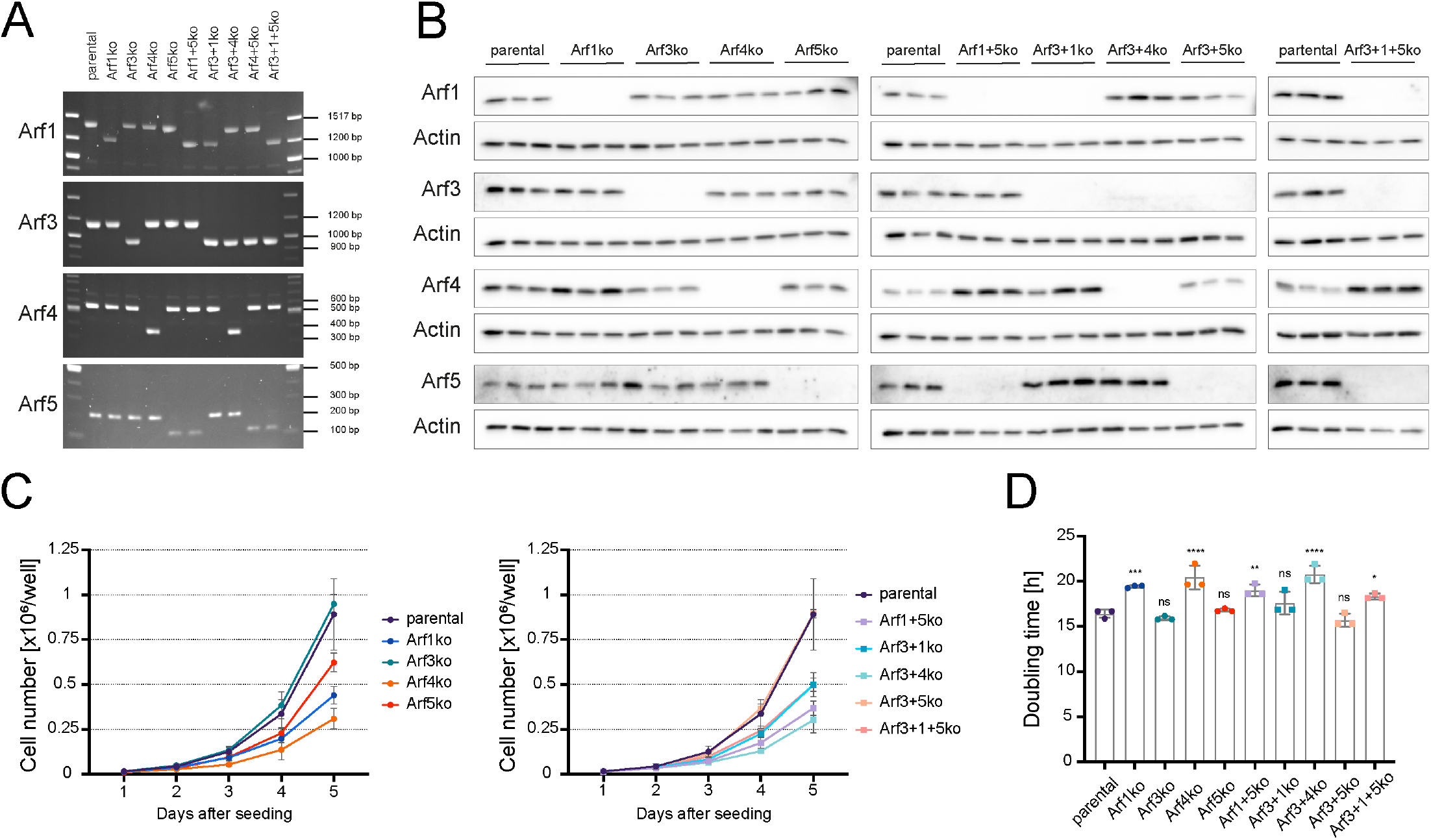
Characterization of viable Arf knockout cell lines. (A) Arf knockout HeLaα cell lines were generated by CRISPR/Cas9 as described in Materials and Methods and analyzed for partial deletion of exon 1 by genomic PCR. Uniform shortening of PCR fragments in knockout cell lines indicated successful genomic deletion in all corresponding Arf alleles. (B) The deletion of Arfs on the protein level was verified by immunoblot analysis. For each cell line, three biological replicates were analyzed on the same gel. (C) Growth curves of Arf knockout cell lines were derived from live cell counts obtained in 3 independent experiments. Cells were seeded at the same density and monitored for 5 consecutive days. (D) Doubling times were calculated from the growth curves displayed in C. Statistical significance was calculated using the unpaired one-way ANOVA vs. parental (ns, not significant, p > 0.05; * p < 0.05; ** p < 0.01; *** p < 0.001; **** p < 0.0001).

To test the impact of Arf deletions on cell growth, the doubling time was determined from growth curves based on live cell counts (Figure 1C and D). The growth rates of Arf3ko, Arf5ko and Arf3+5ko cell lines were comparable to parental HeLaα cells. The other cell lines lacking Arf1 or Arf4, either alone or in combination with another Arf, grew more slowly, resulting in an increase in doubling time of 13% (± 4%) for cells without Arf1 and of 26% (± 1%) for Arf4 deleted cell lines.

We conclude that no single Arf is required for cell survival. This is also true for several Arf knockout combinations, with the exception of Arf4 in combination with Arf1 or Arf5. Simultaneous deletion of these Arfs appears to be lethal, since the respective knockout cell lines could not be generated. Remarkably, Arf4 alone is sufficient for cell viability in the absence of all other class I and II Arfs. This highlights that class I Arfs are not essential.

### Knockout of specific Arfs affects Golgi morphology and steady-state localization of coats

First, we analyzed the impact of Arf deletions on the morphology of the Golgi complex and the steady-state localization of associated coat components by confocal immunofluorescence microscopy. Upon staining for the *cis*-Golgi golgin GM130, the Golgi appeared as a perinuclear compact tangle of ribbons in parental HeLaαcells (Figure 2A). In all cell lines lacking Arf1 (Arf1ko, Arf1+5ko, Arf3+1ko, Arf3+1+5ko), however, this shape was altered to a more diffuse and less compact pattern that appeared swollen and enlarged. Similar changes were observed also for the staining of the TGN marker TGN46, which covered an expanded area in all cell lines lacking Arf1 compared to parental cells (Figure 2B and C). In addition, the mean intensity of the TGN46 signal appeared to be lower. No alteration was observed by eye in Golgi morphology in the other Arf knockout cell lines.

**Figure 2:**
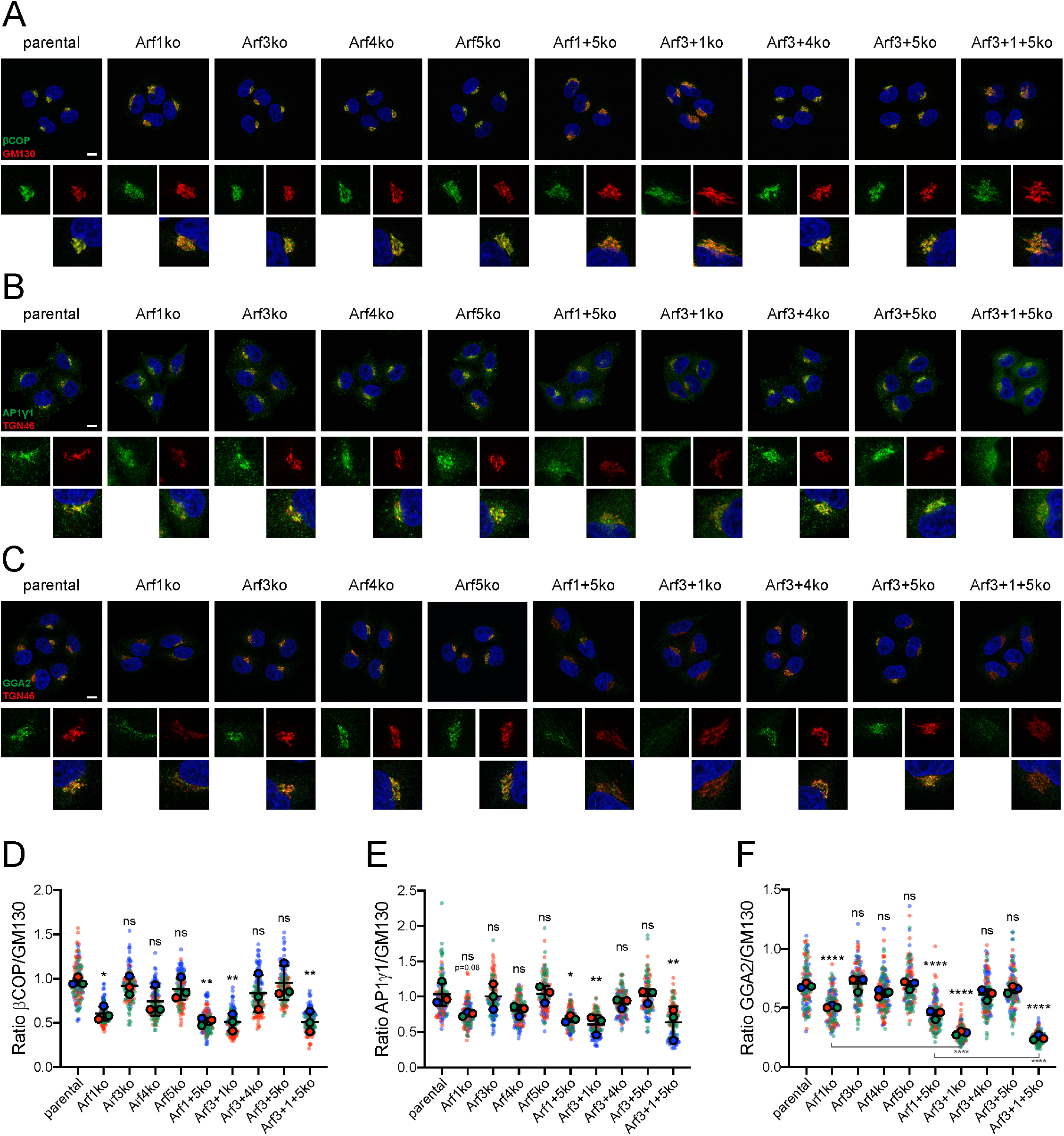
Immunostaining for the Golgi and associated coat proteins. All Arf knockout cell lines, as well as parental HeLaα cells, were immunostained for the marker proteins GM130 (A) and TGN46 (B and C) to examine Golgi morphology (red). Steady-state localization of the coat components βCOP (A), AP1γ1 (B) and GGA2 (C) was determined by co-immunostaining (green). DAPI signal is shown in blue. Lower panels show magnification (5x) of image sections. Scale bar, 10 μm. Perinuclear localization of βCOP (D), AP1γ1 (E) and GGA2 (F) was quantified in relation to the co-stained GM130. Average values obtained from 3 independent experiments are shown in intense colours and the underlying values from individual cells are shown semi-transparent in the background. Statistical significance was calculated using the unpaired one-way ANOVA vs. parental, unless indicated otherwise (ns, not significant, p > 0.05; * p < 0.05; ** p < 0.01; *** p < 0.001; **** p < 0.0001).

By double-staining, we simultaneously examined the steady-state localization of three Golgi-associated, Arf-dependent coats, COPI, AP1 and GGA2 (Figure 2A-C, respectively). The signal for βCOP, AP1γ1, and GGA2 mostly colocalized with the Golgi markers. Additional puncta were detected in the cell periphery, most strongly for AP1γ1, well known to also localize to early endosomes, more weakly for GGA2 and βCOP. For all three coat components, the signal appeared to be reduced at the Golgi in all Arf1 knockout cells lines (Arf1ko, Arf1+5ko, Arf3+1ko, Arf3+1+5ko) compared to parental HeLaγcells. Moreover, the endosomal AP1#x03B3;1 was almost lost. No change was observed in the other Arf knockout cell lines.

To quantify the change in coat proteins at the Golgi, we co-stained the cells for GM130 (Figure 2A and Suppl. Figure S1) to define the region of interest and used its mean fluorescence intensity, which appeared to be constant between cell lines, for normalization. The results forβCOP and AP1γ1 confirmed a reduction of the coat signal in the Golgi region by an average of 40-50% and 30-40%, respectively, in all Arf1 knockout cell lines compared to parental cells (Figure 2D and E). For cell lines lacking Arf4 (Arf4ko, Arf3+4ko), a slight, although not significant decrease of Golgi-localizedβCOP and AP1γ1 was observed.

GGA2 at the Golgi was also reduced like AP1γ1 in all Arf1 deleted cell lines (Figure 2F). Remarkably, the additional absence of Arf3 in Arf3+1ko and Arf3+1+5ko cells even enhanced the loss of GGA2 at the Golgi compared to Arf1ko and Arf1+5ko cells from ∼30% to ∼60%. Again, a slight, non-significant reduction of GGA2 was detected in cells lacking Arf4.

The results suggest that normally Arf1 is the most important mediator of Golgi recruitment of the coats tested.

### Golgi volume is increased and 3D morphology altered in cells deleted for Arf1 or Arf4

To substantiate the initial observation of differences in Golgi size, we generated serial z-stack images of cells immunostained for GM130. In maximum intensity projections, the Golgi complex again appeared enlarged and more diffuse in cells lacking Arf1 (Figure 3A). Subtle changes were also suspected for cell lines lacking Arf4, as the GM130-positive ribbons appeared to be more densely packed.

**Figure 3:**
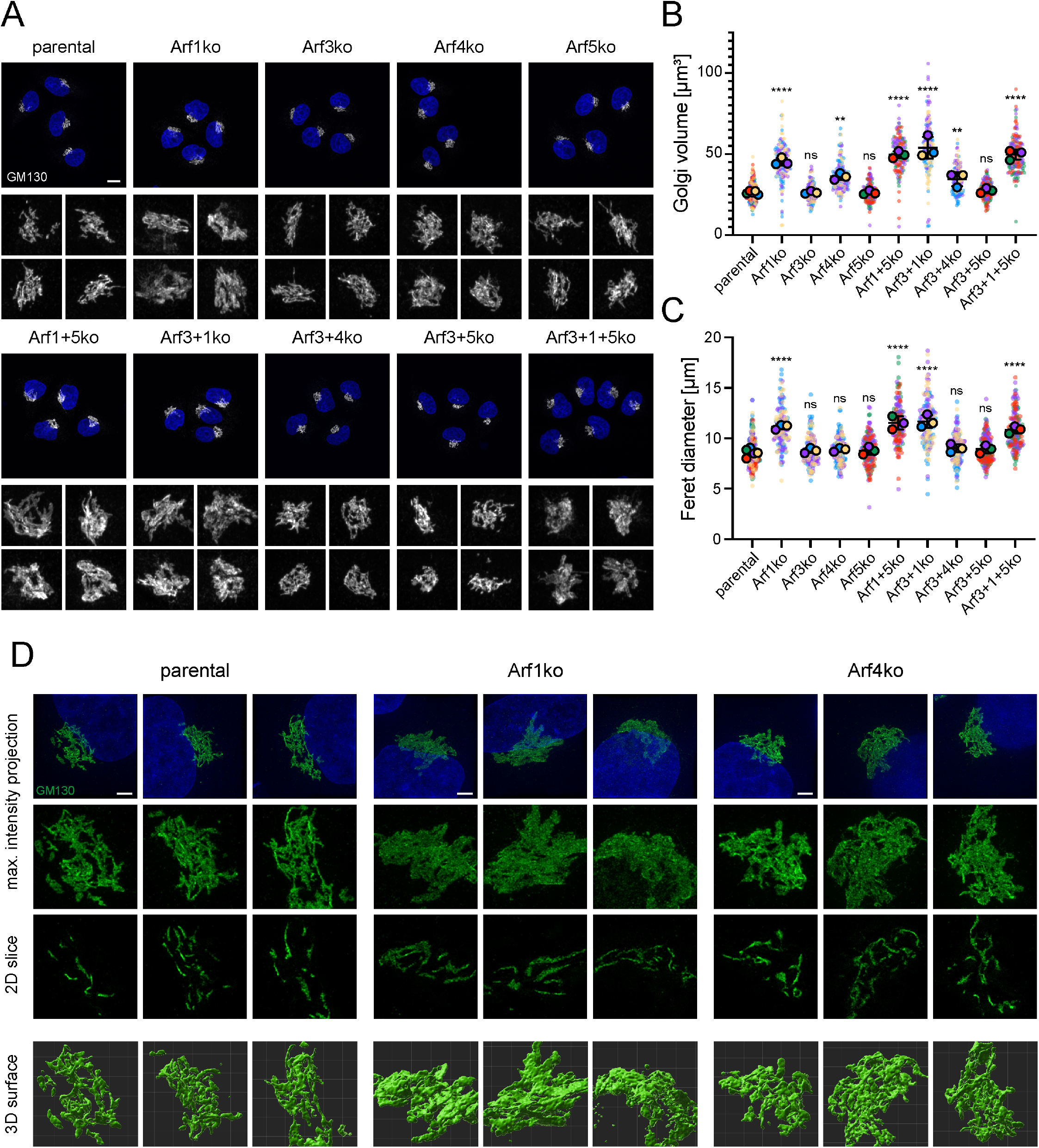
Impact of Arf knockouts on Golgi morphology. (A) Maximum intensity projections were generated from serial confocal z-stack images of GM130-labeled Golgi. Lower panels show a magnified view (7.5x) of the Golgi complexes. Scale bars, 10 μm. Volume (B) and Feret diameter (C) of individual Golgi complexes were determined in 3 independent experiments using z-stack images described in A (on average 50 Golgi per cell line and experiment). Average values are shown in intense colours and the underlying values from individual Golgi are shown semi-transparent in the background. Statistical significance was calculated using the unpaired one-way ANOVA vs. parental (ns, not significant, p > 0.05; * p < 0.05; ** p < 0.01; *** p < 0.001; **** p < 0.0001). (D) GM130 stained Golgi were imaged by superresolution microscopy and results are displayed as maximum intensity projections, tomographic 2D slices and 3D reconstructed surfaces. DAPI is shown in blue. Lower panels show a magnified (2x) image section. Scale bar, 3 μm.

Golgi volume and Feret diameter (the largest distance between two contour voxels) were measured after 3D reconstruction and confirmed the visual evaluation (Figure 3B and C). All cell lines deleted for Arf1 displayed a significant increase in Golgi volume (on average 1.9-fold) and in the Feret diameter (on average 1.3-fold). In cell lines lacking Arf4, the Golgi volume was also significantly increased (on average 1.3-fold), whereas the Feret diameter remained at the levels of parental HeLaα cells. The other knockout cell lines (Arf3ko, Arf5ko, Arf3+5ko) did not exhibit a change in either parameter.

How these changes manifest themselves in the 3D structure of the Golgi, was addressed by superresolution microscopy (3D structured illumination microscopy; Figure 3D). In maximum intensity projections of 3D SIM z-stacks, the GM130 labeled Golgi of parental HeLaα cells presented itself as a network of ribbons. In Arf1ko cells, this pattern was altered and appeared less defined and more uniformly distributed. Arf4ko Golgi displayed an intermediate phenotype. Tomographic 2D slices showed individual ribbons that were increased in the diameter in Arf1ko cells. In Arf4ko Golgi, the ribbons seemed to be more intertwined than in parental cells.

Golgi surface reconstruction showed a marked difference between a tangle of tubular structures in parental cells, clusters of more planar sheets in Arf1ko cells, and more densely packed Golgi ribbons in Arf4 knockout cells. Taken together, the 3D analysis of Golgi in knockout cell lines confirmed an increase in Golgi volume for Arf1ko and Arf4ko cell lines and linked it to a broadening or higher number of Golgi ribbons, respectively.

### Morphological changes in the ultrastructure of Arf knockout cells

We further analyzed the ultrastructure of the Golgi by thin-section transmission electron microscopy (Figure 4A). The only difference in the appearance of Golgi structures was an increase in the length of individual stacks in cell lines lacking Arf1 compared to parental HeLaα cells. Quantitation revealed a significant increase in stack length of ∼65% in Arf1ko cells, while stack thickness and the number of cisternae per stack remained unchanged (Figure 4B–D). This is consistent with the increased ribbon diameter observed by superresolution microscopy. No indication of morphological changes in Golgi structure was seen in other cell lines.

**Figure 4:**
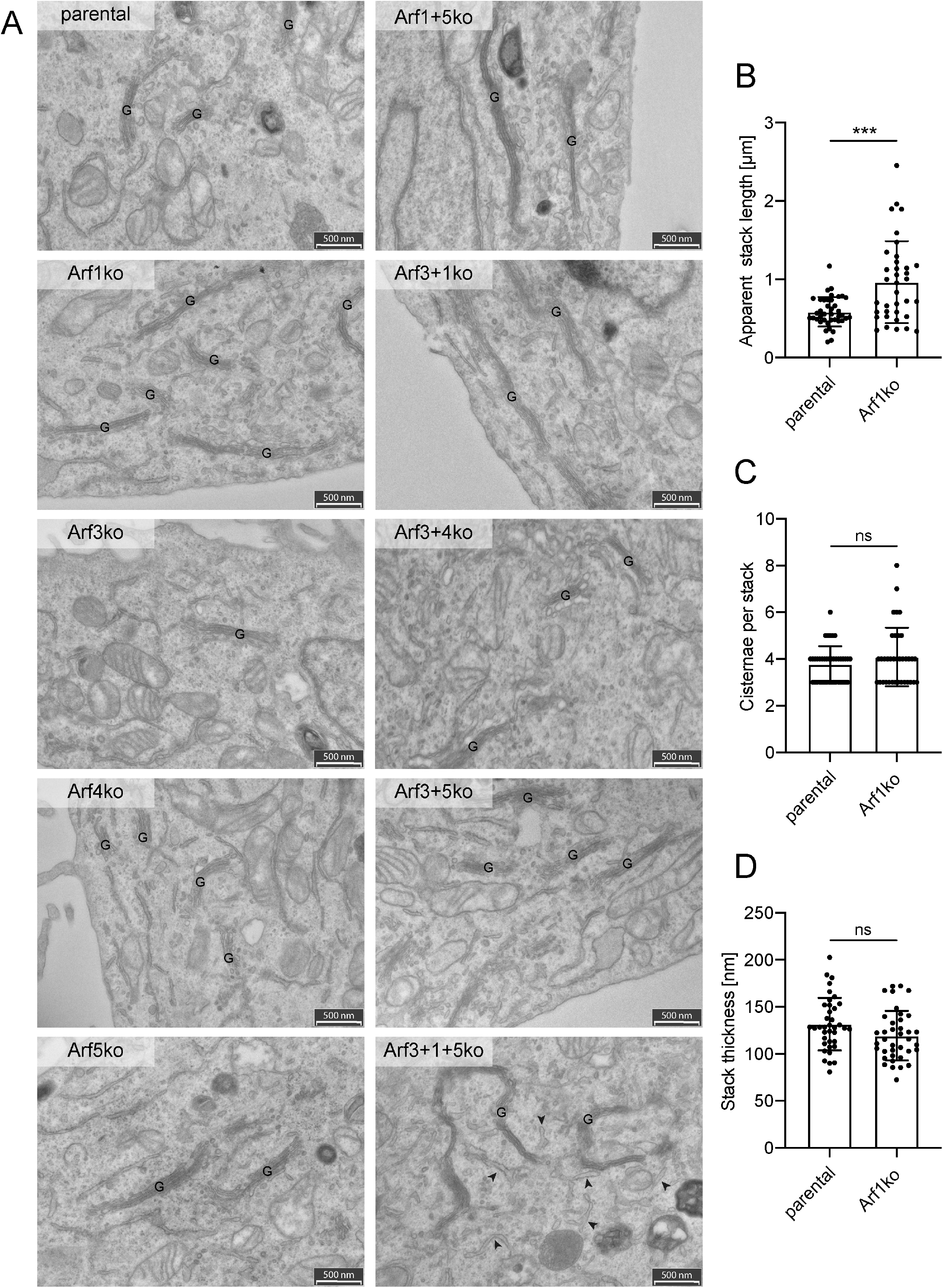
Analysis of the ultrastructure of the Golgi. (A) Thin section transmission electron microscopy (TEM) was performed to examine the ultrastructure of cellular compartments in all Arf knockout cell lines. The field of view was chosen to include Golgi stacks in the acquired images. G, Golgi stack; Arrowheads indicate dilated ER; Scale bar, 500 nm. (B) Individual Golgi stacks were quantified for their apparent length, number of cisternae per stack and thickness in electron microscopy images of HeLaα and Arf1ko cells. Approximately 40 Golgi stacks were quantified per cell line. Statistical significance was calculated using the unpaired, two-tailed t-test (ns, not significant, p > 0.05; * p < 0.05; ** p < 0.01; *** p < 0.001; **** p < 0.0001).

Interestingly, in the triple-knockout cell line (Arf3+1+5ko) the tubular elements of the ER, as identified by bound ribosomes, appeared frequently dilated (Figure 4A, arrowheads).

### Knockout of Arf4 causes defective retrieval of ER resident proteins

To assess the functionality of the secretory pathway upon deletion of Arfs, we analyzed total secretion by visualizing secreted proteins collected from the media by SDS-gel electrophoresis and Coomassie staining (Figure 5A). Strikingly, in the media of Arf4 deleted cell lines, additional bands were detected. For identification of these additionally secreted proteins, media was collected from Arf4ko, Arf3+4ko, and parental HeLaα cells and analyzed by mass spectrometry. This approach identified 75 and 87 proteins to be significantly up-regulated (as defined by fold change >2, q-value <0.01) in the secretomes of Arf4ko and Arf3+4ko cell lines, respectively (Figure 5B, Suppl. Table S2 and S3). Seventy of these were shared between the two knockout cell lines. Among the top hits, we found ER chaperones, such as BiP, calreticulin, and GRP94, peptidyl-prolyl cis/trans isomerases, and protein disulfide isomerases. Gene ontology term (GOterm) enrichment analysis identified the ER as the main compartment of origin (Suppl. Table S4 and 5). Secretion of BiP and calreticulin was verified by immunoblot analysis (Figure 5C). Both chaperones were found to be strongly secreted specifically in the two cell lines lacking Arf4.

**Figure 5:**
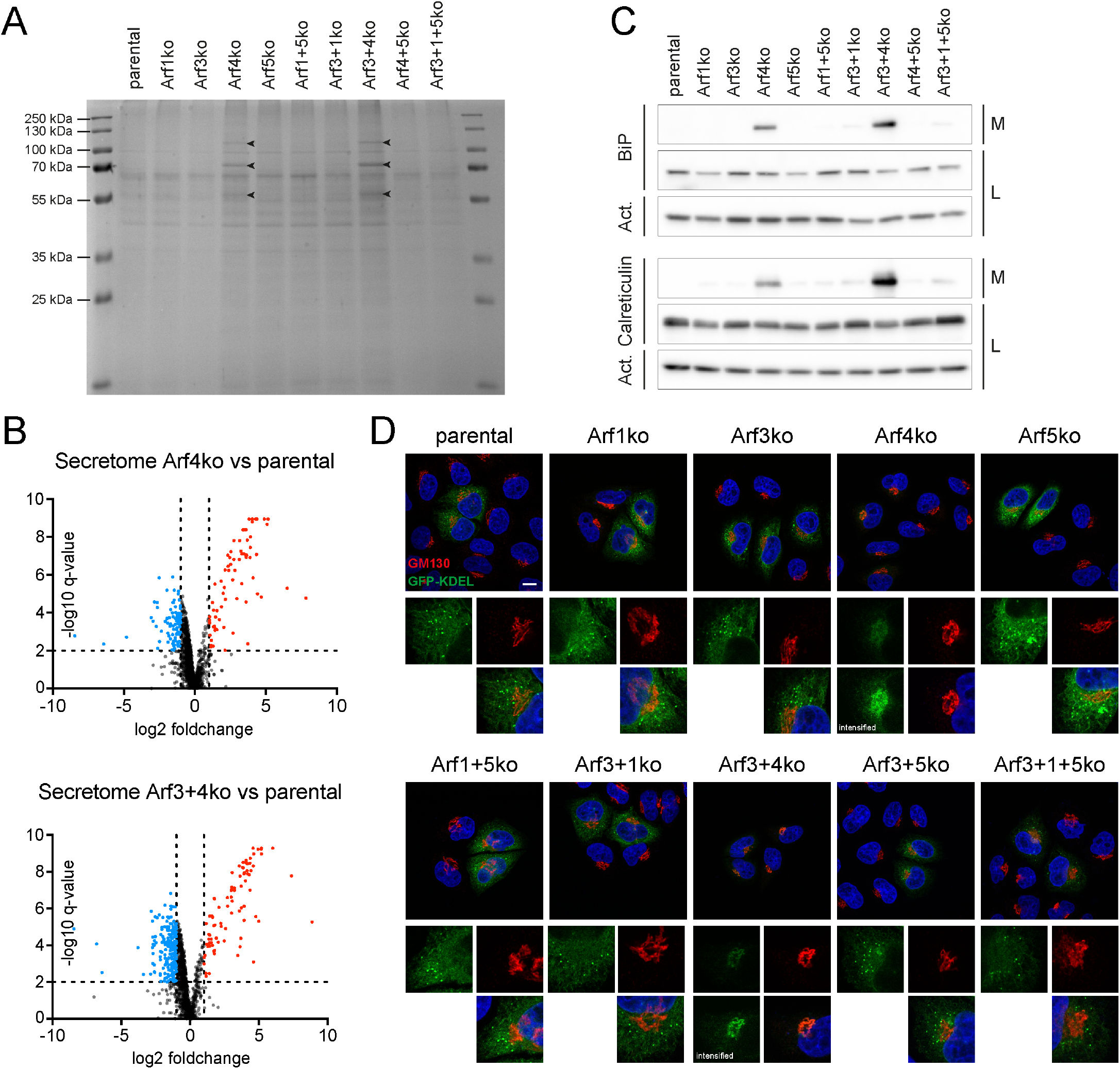
Examination of total secretion and KDEL-based retrieval of ER resident proteins. (A) Secreted proteins collected from the media of Arf knockout cell lines were visualized on Coomassie stained SDS-gels. Arrowheads indicate additional bands. (B) Vulcano plot of approximately 1400 secreted proteins, which were detected by mass spectrometry in the media of parental HeLaα, Arf4ko and Arf3+4ko cells. Proteins that were found to be enriched in the knockout compared to parental samples were marked in red (foldchange > 2, q-value <0.01) and depleted ones were marked in blue (foldchange < 0.5, q-value <0.01). Analysis was done in biological quintuplicates. Dotted lines indicate the significance thresholds. (D) Media collected from the indicated cell lines(M) and the corresponding cell lysates (L) were probed for the presence of the two ER-chaperones BiP and calreticulin by immunoblotting. Actin (Act.) served as a loading control. (E) Steady-state localization of transiently expressed signal sequence-GFP-KDEL (green) co-immunostained with GM130 (red). DAPI signal is shown in blue. Lower panels show magnified image sections (5x). Scale bar, 10 μm.

Aberrant secretion of ER resident proteins in Arf4 deleted cells indicates a defect in their retrieval from the Golgi back to the ER. The most prominent mechanism is retrograde transport by the KDEL receptors (Lewis and Pelham 1990a, Hsu et al. 1992, Lewis and Pelham 1992b, Raykhel et al. 2007). Indeed, ∼30% of the proteins, whose secretion was increased in Arf4 knockout cells, contain a KDEL motif at their C-terminus or a variant thereof according to the PROSITE consensus sequence [KRHQSA]-[DENQ]-E-L> (entry PDOC00014). However, this might be an underestimation as studies suggested that not all motifs recognized by the KDEL-receptors are included in this consensus pattern (Raykhel et al. 2007).

Functionality of KDEL-mediated retrieval in Arf knockout cell lines was tested by transient expression of a signal sequence-GFP-KDEL (GFP-KDEL) construct and examination of its steady-state localization (Figure 5D). In parental HeLaα cells, the GFP signal was visible in the reticular pattern of the ER and a number of puncta. Exclusively in cells lacking Arf4, the GFP-KDEL-positive puncta were completely lost and the reticular staining reduced. Instead, a perinuclear accumulation colocalizing with GM130 was detected consistent with a defective retrieval from the Golgi back to the ER in the absence of Arf4. Defective retrieval and aberrant secretion of proteins with a KDEL motif in cells lacking Arf4 indicates an important role for Arf4 in retrograde transport from the Golgi to the ER.

### Knockout phenotypes are rescued by overexpressing Arfs of the same class

To test the specificity of the observed phenotype for the deleted Arf by rescue experiments, we generated Arf1ko cells stably re-expressing Arf1 or overexpressing the other class I family member Arf3, and Arf4ko cells stably re-expressing Arf4 or overexpressing the other class II member Arf5. As controls, Arf1–5 were stably overexpressed in parental HeLaα cells. The stable cell lines were generated by lentiviral transduction, introducing untagged versions of the corresponding Arfs. Clonal cell lines with a comparable expression level in parental and knockout background were selected based on immunoblot analysis (Figure 6A). Compared to the endogenous Arf levels in parental HeLaα cells, Arf1 and Arf4 were moderately and Arf3 and Arf5, which have low endogenous levels, were highly overexpressed.

**Figure 6:**
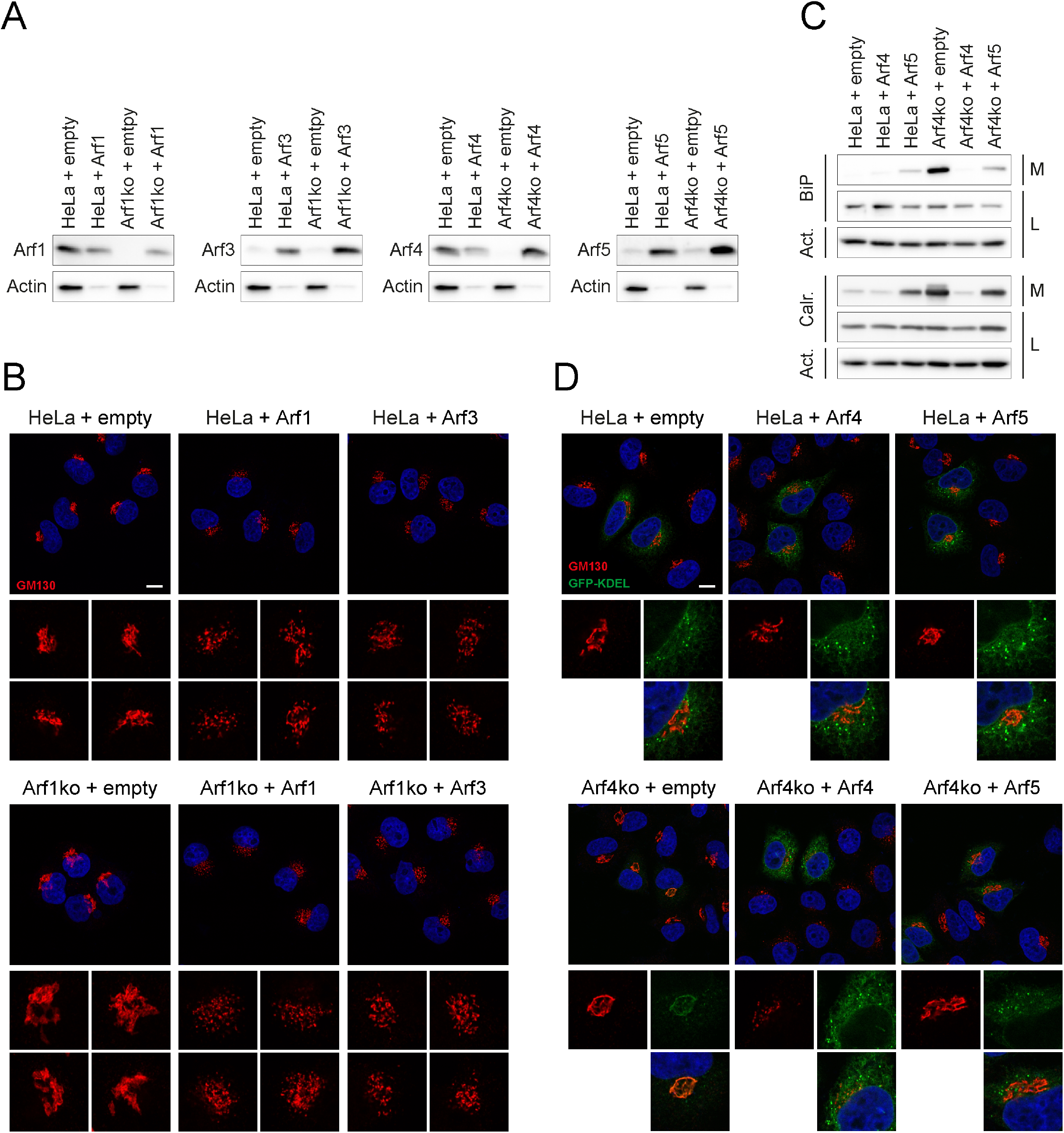
Rescue experiments overexpressing Arfs of the same class. Based on parental HeLaα, Arf1ko and Arf4ko cells, stable cell lines were generated by lentiviral transduction, either expressing the empty vector (+empty) as a control or overexpressing the indicated, untagged Arf. (A) Expression levels of Arfs in clonal cell lines were analyzed by immunoblotting. Twenty times less lysate was loaded for Arf overexpressing cell lines. Actin served as a loading control. (B) Parental HeLaα and Arf1ko cell lines transduced with the empty vector or overexpressing class I Arfs were immunostained for GM130 to examine Golgi morphology (red). Magnification of image sections (5x) are shown in the lower panels. Scalebar, 10 μm. (C) Immunoblot analysis probing the media (M) and cell lysates (L) of parental HeLaα and Arf4ko cell lines transduced with the empty vector or overexpressing class II Arfs for the chaperones BiP and calreticulin (Calr.). Actin (Act.) served as a loading control. (D) Immunofluorescent microscopy revealed the steady-state localization of transiently expressed signal sequence-GFP-KDEL (green) co-immunostained with GM130 (red) in parental HeLa α and Arf4ko cell lines transduced with the empty vector or overexpressing Arf4 or Arf5. DAPI signal is shown in blue. Lower panels show magnified image sections (5x). Scale bar, 10 μm.

The cell lines overexpressing class I Arfs were examined for their Golgi morphology by immunofluorescence staining of GM130 (Figure 6B). In parental HeLaα cells, overexpression of Arf1 or Arf3 showed an opposite effect of Arf1 deletion: the Golgi condensed onto individual filaments and puncta. The same phenotype was also obtained in Arf1ko cells, overriding the knockout phenotype.

In the cell lines overexpressing class II Arfs, we assessed the restoration of ER protein retrieval (Figure 6C and D). As demonstrated by immunoblot analysis, aberrant secretion of BiP and calreticulin in Arf4ko cells was completely reverted to background levels upon overexpression of Arf4 and also partially upon overexpression of Arf5 (Figure 6C). Overexpression of Arf4 in wild-type HeLaα cells had no effect. Remarkably, Arf5 overexpression in HeLaα cells caused significant secretion of these ER chaperones. We speculate that Arf5 cannot completely substitute for Arf4 with respect to ER protein retrieval and, if Arf4 is present, overexpressed Arf5 competes with Arf4, but is less productive.

Immunofluorescence microscopy of GFP-KDEL confirmed these observations by full rescue of ER staining with punctate accumulations in Arf4ko cells overexpressing Arf4 and a slightly weaker rescue by Arf5 overexpression (Figure 6D). Moreover, in cells overexpressing Arf4, the GM130-stained Golgi appeared disrupted. This was not observed in Arf5 overexpressing cells.

Reversal of the deletion phenotypes by re-expression confirms that they were caused by the particular knockout. In both cases, the overexpression of the other member of the same class produced similar results, although to different extents. This suggests at least partial redundancy between the Arfs of the same class.

## Discussion

### No single Arf is essential and Arf4 is sufficient for cell survival

Arf GTPases are important regulators of a range of cellular processes, especially within the secretory pathway. For this reason, their misregulation is associated with diverse human diseases, including cancer (reviewed in Casalou et al. 2016, Sztul et al. 2019). Hence, it is important to decipher and understand shared and specific functions of individual Arfs. Their identification has been complicated, for instance, by apparent functional redundancy, shared GEFs and GAPs, a complex interactome, as well as technical difficulties (Sztul et al. 2019). Previous analyses employed shRNA- or siRNA-mediated knockdowns, potentially prone to incomplete depletion and off-target effects, or (over)expression of mutant Arfs. Dominant-active (GTPase-deficient) or dominant-negative (GDP-locked) mutants may interfere with the activity of other Arfs by outcompeting them or by blocking shared GEFs. Furthermore, epitope-tagging was shown to change the Arfs’ properties in subtle ways (Jian et al. 2010).

Our approach to gain insight into Arf functions was based on genomic deletions of individual Arfs and Arf combinations and subsequent observation of defects in cell growth, Golgi morphology and coat recruitment to the Golgi. We successfully generated nine genomic Arf knockout cell lines, comprising four single-knockouts of each human class I (Arf1, Arf3) and class II (Arf4, Arf5) Arf, four double- and one triple-knockout combination. These cells are permanently depleted for the respective Arfs, which may lead to compensatory effects. In this line, we only observed an upregulation of Arf4 in all cell lines lacking Arf1 (Fig 1B), but no further changes in Arf expression levels.

In agreement with knockdown based studies, we found that cultured cells can cope rather well with the loss of any single Arf and even certain combinations. However, the single-knockout of Arf1 or Arf4 exhibited significant and distinct phenotypes, for example, either deletion reduced the growth rate. The combined Arf1+4 double-knockout was not viable. Interestingly, the same is true for simultaneous deletion of both class II Arfs (4+5), whereas cells knocked out for both class I Arfs (1+3) grew as well as Arf1ko cells. In addition, Arf4 was found to be able to sustain all essential Arf functions alone in cultured cells lacking the other three Arfs.

### Deletion of Arf1 increases Golgi volume and reduces coat recruitment

The most striking phenotype of cells deleted for Arf1, alone or in combination with Arf3 and/or Arf5, was an enlargement of the Golgi, apparent in a more expanded GM130 staining, a higher volume in 3D Golgi reconstructions, and longer Golgi stack cross sections in electron microscopy. Furthermore, the Arf1ko cell line displayed reduced steady-state levels of COPI, AP1, and GGA2 vesicle coat components at the Golgi, indicating reduced formation of COPI and AP1/GGA/clathrin transport vesicles originating from the Golgi in the retrograde and anterograde direction, respectively. Consequential reduction of cargo and membrane flow out of the Golgi in both directions and a resulting imbalance of influx and efflux might explain the increased Golgi size, as proposed by Sengupta and Linstedt (2011).

An increase in Golgi volume has also been observed physiologically upon an increased demand for cargo transport, processing, and sorting at the Golgi (Sengupta and Linstedt 2011). As cells grow during interphase, increased cargo load causes a Golgi volume increase. Golgi growth was shown in HeLa cells to occur by cisternal elongation of existing Golgi stacks rather than by addition of new stacks (Sin and Harrison 2016). Thus, the observed length increase in Golgi stacks of our Arf1 knockout cells is likely to results from increased cargo content due to reduced export rates. Taken together, Arf1 appears to be the major mediator of vesicle traffic originating from the Golgi.

### Knockout of Arf4 specifically disrupts retrieval of ER proteins

The deletion of Arf4, alone or in combination with Arf3, caused a slight increase in Golgi volume and a reproducible, but not statistically significant decrease of coat components at the Golgi. Arf4 thus appears to contribute to the recruitment of COPI, AP1 and GGA2 and the consequential mild reduction of Golgi exit might explain the observed slight increase in Golgi volume in Arf4ko cell lines.

Most strikingly, however, Arf4ko cells manifest an aberrant secretion of ER resident proteins that are normally retrieved from the Golgi back to the ER by the KDEL receptors. The phenotype resembles the one reported for knockdowns of multiple KDEL receptors (Li et al. 2015). However, this defect upon Arf4 deletion cannot be simply linked to a general defect in retrograde transport due to reduced COPI recruitment, since this is more strongly observed in Arf1ko cells without causing ER protein secretion. A more specific function in ER protein retrieval must therefore be defective in Arf4ko cells.

It has previously been shown that in addition to Arf1 also the class II Arfs, but not Arf3, can competitively support COPI vesicle formation (Popoff et al. 2011). For COPI, it has been shown that two paralogs of γ-COP (γ1/γ2) and of ζ-COP (ζ1/ζ2) can form three distinct COPI complexes (γ1ζ1, γ1ζ2, or γ2ζ1) with potentially different functions (Moelleken et al. 2007, Wegmann et al. 2004). Furthermore, Scyl1, a member of the Scy1-like family of catalytically inactive protein kinases, was identified as an interactor of COPI at the *cis*-Golgi and ERGIC that causes reduced retrograde traffic of the KDEL receptors when inactivated (Burman et al. 2008, Burman et al. 2010). Subsequently, Scyl1 was shown to bind to class II Arfs, preferentially Arf4, and to interact directly with the COPI subunit γ2 (Hamlin et al. 2014). This interaction was recently shown to depend on arginine methylation of Scyl1 by PRMT1 (Amano et al. 2020). A tripartite Scyl1–Arf4–γ2ζ1-COP complex thus was proposed to specifically mediate KDEL receptor traffic. This mechanism may thus account for the specific phenotype we observe upon Arf4 deletion.

However, in other studies Scyl1 was also reported to bind preferentially to Arf1 and to GORAB, a protein associated with gerodermia osteodysplastica, at the *trans*-Golgi to promote COPI recruitment (Witkos et al. 2019). Pathogenic GORAB mutations cause impairment of COPI-mediated retrieval of *trans*-Golgi enzymes and a deficit in glycosylation of secretory proteins. Based on their results, the authors suggest that there might be two separate pools of Scyl1, a GORAB-dependent one at the *trans*-Golgi and a pool at the *cis*-Golgi/ERGIC for distinct COPI functions.

The situation is further complicated by the recent finding that a mutation in γ1-COP (K652E), shown to cause defective humoral and cellular immunity, disrupted KDEL receptor binding to COPI, thus affecting KDEL receptor localization, increasing ER stress in activated T and B cells and apoptosis in activated T cells (Bainter et al. 2021). Thus, other, γ1-containing COPI complexes also appear to contribute to KDEL receptor sorting in these cells. How Arf4 specifically mediates KDEL receptor retrieval is therefore not entirely clear yet.

### Arf knockout combinations

In the majority of viable Arf double- or triple-knockout cell lines, no additional phenotypes were detected beyond those of Arf1 or Arf4 single deletion regarding Golgi size and morphology, coat recruitment, or secretion of ER resident proteins. However, in Arf3+1ko cells, the loss of GGA2 from the Golgi was more pronounced than in Arf1ko cells. This suggests a functional overlap of Arf1 and Arf3 in the recruitment of GGA2, which is consistent with Arf3’s known preferential localization to the TGN and activation by the *trans*-Golgi GEF BIG1 (Manolea et al. 2010) and its ability to bind GGAs (Boman et al. 2000). The only exception is the additional observation of a dilated ER in the Arf3+1+5 knockout cell line.

Previous knockdown-based studies reported phenotypes only upon simultaneous silencing of two Arfs (Volpicelli-Daley et al. 2005, Kondo et al. 2012, Nakai et al. 2013). KDEL-receptor localization, for example, was described to be enhanced at the Golgi upon double knockdown of Arf3+Arf4 (Volpicelli-Daley et al. 2005) and of Arf4+Arf5 (Volpicelli-Daley et al. 2005, Li et al. 2015). Our results attribute this phenotype solely to the deletion of Arf4. The same applies to a slight compaction of the Golgi observed upon knockdown of Arf4+Arf5 (Nakai et al. 2013).

In other cases, the reported phenotype, for instance the peripheral βCOP puncta observed upon knockdown of Arf1+Arf3 and Arf1+Arf5 (Volpicelli-Daley et al. 2005, Kondo et al. 2012), cannot be rationalized by our knockouts. Knockdown of Arf combinations for which no knockout cell lines could be generated could provide information on the defects that lead to growth arrest or cell death. In this line, the simultaneous knockdown of Arf1+Arf4 severely impacted Golgi morphology and AP1 and COPI localization (Volpicelli-Daley et al. 2005; Nakai et al. 2013).

In the present study, we established Arf knockout cell lines as tools to study shared and specific functions of Arfs at the Golgi. Of course, these cell lines offer themselves to analyze Arf-dependent processes that do not require specialized cell types.

## Material and Methods

### Cell culture

Helaγ cells were grown in DMEM (high glucose; Sigma-Aldrich) with 10% fetal calf serum (FCS premium, S. Amer. Orig, VWR), 2 mM L-glutamine, 100 units/ml penicillin G, and 100 ng/ml streptomycin at 37°C and 7.5% CO2.

### HeLaα knockout cell lines

Two guide RNAs (gRNAs) were designed for each targeted Arf gene to facilitate genomic deletion of exon 1 using several online tools (Zhang lab; CRISPOR, Concordet and Haeussler 2018; Deskgen, Desktop Genetics), as listed in Suppl. Table S1. gRNAs were cloned in the pSpCas9(BB)-2A-GFP (a gift from Feng Zhang, Addgene plasmid #48138), and pSpCas9(BB)-2A-mCherry, which was derived the former by exchanging GFP to mCherry. Target cells were transfected simultaneously with the corresponding plasmids using jetPRIME (Polyplus Transfection). After 24 h, double-fluorescent cells were selected by FACS (FACS AriaIII, BD Biosciences). After 7 days, double-negative cells were selected by FACS and single cells sorted into 96-well plates with growth medium containing 10% conditioned medium. After ∼14 days, clonal cell lines were expanded and analyzed.

### Genomic PCR

Trypsinized cells were pelleted, resuspended in 10 mM Tris (pH 8.7), heated at 95°C for 10 min, incubated with proteinase K (0.5 μg/μl) for 20 min at 37°C, inactivated at 95°C for 15 min, and used as DNA template for PCR with Phusion Polymerase (NEB) or Q5 Polymerase (NEB, for Arf5 knockouts), following the manufacturer’s protocol for high GC content. Primers designed to amplify the genomic region surrounding the site of deletion are listed in Suppl. Table S1.

### Immunoblot analysis

Cell lysates were separated by SDS-gel electrophoresis (15% polyacrylamide for Arfs) and transferred to Immobilon-P PDVF membranes (Millipore). Membranes were blocked with TBST (20 mM Tris, 150 mM NaCl, pH 7.6, 0.1% Tween20) with 3% non-fat dry milk for 1 h and incubated with primary antibody in TBST with 1% milk over night at 4°C: anti-Arf1 (1:2500, Abnova MAB10011), anti-Arf3 (1:1000, BD Bioscience 610784), anti-Arf4 (1:1000, Proteintech 11673-1-AP), anti-Arf5 (1:750, Abnova H00000381-M01), anti-actin (1:100000, Sigma-Aldrich MAB1501), anti-calreticulin (1:2500, Proteintech 27298-1-AP), anti-Grp78/Bip (1:10000, Genetex GTX113340-10). After washing, the membranes were incubated with HRP-conjugated secondary antibody (1:10000; anti-rabbit, Sigma-Aldrich A0545; anti-mouse, Sigma-Aldrich A0168) in TBST with 1% milk. Chemiluminescence signals were detected using Immobilon Western HRP Substrate (Millipore) or Radiance Plus (Azure Biosystems) and imaged using a FusionFX (Vilber Lourmat).

### Growth assay

Cells were seeded in 12-well plates at a density of 5500 cells/well, which was confirmed by re-counting. Every 24 h for five consecutive days, cells from one well for each cell line were trypsinized, resuspended in PBS, and counted. Doubling times were estimated by exponential fitting of the growth curves.

### Immunofluorescence staining

Cells were grown on glass coverslips for one day, then fixed with 3% PFA for 10 min, quenched with 50 mM NH_4_Cl in PBS for 5 min, permeabilized with 0.2% TritonX-100 in PBS for 10 min, blocked with 1% BSA in PBS for 1 h, and incubated with primary antibodies diluted in 1% BSA in PBS for 1 h: Anti-AP1γ1 (1:1000, self-made from hybridoma cells), anti-γCOP (1:500, CM1, hybridoma supernatant, gift from Dr. Felix Wieland, Heidelberg University), anti-GGA2 (1:500, BD Bioscience 612613), anti-GM130 (1:1000, Cell Signaling 12480S), and anti-TGN46 (1:1000, BioRad AHP500G). Samples were washed and incubated with fluorescent secondary antibodies diluted in 1% BSA in PBS for 1 h (1:400; anti-mouse-Alexa488, Invitrogen A21202; anti-rabbit-Alexa568, Invitrogen A10042; anti-sheep-Cy3, Jackson ImmunoResearch 713-165-147). Coverslips were mounted in FluoromountG (SouthernBiotech) supplemented with 0.5 ng/ml DAPI (Sigma-Aldrich) and stored in the dark at 4°C. For localization of GFP-KDEL, cells were grown on coverslips for one day, transfected with the pcDNA3-ss-GFP-KDEL, and fixed and stained a day later.

### Confocal microscopy and quantitation of coat localization and Golgi volume

Images were acquired using a LSM700 Upright confocal laser-scanning microscope with the Zen 2010 software (Zeiss) equipped with a Plan-Apochromat 63×/1.4 oil-immersion objective lens and two photomultiplier tubes. Imaging parameters were kept constant throughout each experiment. For quantitation of coat proteins at the Golgi, the GM130-stained area was selected in Fiji using the freehand tool and the mean fluorescence intensity was measured for GM130 and the coat protein. To measure Golgi volume, z-stacks at 0.13 μm per slice were acquired and analyzed in Fiji using the “3DGolgiCharacterization” script (DOI: 10.5281/zenodo.4068393).

### Super-resolution microscopy

3D structured illumination microscopy (3D-SIM) was performed on a DeltaVision OMX-Blaze V4 system (Cytiva) equipped with solid-state lasers. Images were acquired using a Plan Apo N 60x, 1.42 NA oil immersion objective lens (Olympus), and 4 liquid-cooled sCMOS cameras (pco.edge 5.5, full frame 2560 x 2160; PCO). Exciting light was directed through a movable optical grating to generate a fine-striped interference pattern on the sample plane. The pattern was shifted laterally through five phases and three angular rotations of 60° for each z section. The 405 and 488 nm laser lines were used during acquisition and the optical z-sections were separated by 0.125 μm. For the acquisition at 405 nm, laser power was attenuated to 50% with an exposure time of 40 ms, for 488 nm to 10% and 6ms. Settings were adjusted to achieve optimal intensities of between 5,000 and 8,000 counts in a raw image of 15-bit dynamic range at the lowest laser power possible to minimize photobleaching. Multichannel imaging was achieved through sequential acquisition of wavelengths by separate cameras.

Raw 3D-SIM images were processed and reconstructed using the DeltaVision OMX SoftWoRx software package (v6.1.3, Cytiva). The resulting size of the reconstructed images was of 512 × 512 pixels from an initial set of 256 × 256 raw images. The channels were aligned in the image plane and around the optical axis using predetermined shifts as measured using a target lens and the SoftWoRx alignment tool. The channels were then carefully aligned using alignment parameter from control measurements with 0.5 μm diameter multi-spectral fluorescent beads (Invitrogen, Thermo Fisher Scientific). For visualization of the Golgi surface, we used the surface tool of the Imaris Cell Imaging software (Oxford Instruments).

### Electron microscopy

Cells were fixed in serum-free medium with 2.5% glutaraldehyde and 3% formaldehyde for 2 h at room temperature, washed with PBS and incubated with 2% osmium tetroxide and 1% K-hexacyanoferrate in H_2O_ for 1h at 4°C. After washing with H_2O_, uranyl-acetate (2% in H_2O_) was added and incubated at 4°C overnight. Cells were scraped after washing with H_2O_, pelleted, dehydrated by sequential incubation in 20%, 50%, 70%, 90%, and three times 100% acetone/H_2O_ for 30 min each, infiltrated with EMbed-812 (Electron Microscopy Science) according to the manufacturer’s protocol, and allowed to polymerize for 24 h at 60°C. The embedded cell pellets were cut into 60–70 nm thin sections using an ultramicrotome (UltracutE, Reichert-Jung), collected on carbon-coated Formvar-Ni grids (Electron Microscopy Science), and stained for 10 min in 4% uranyl acetate and 2 min with lead citrate. Images were acquired on a CM100 electron microscope (Philips).

### Analysis of secreted proteins

Cells were washed with PBS and incubated for 1 h with serum-free medium in the incubator. After another wash, secreted proteins were collected in a small volume of serum-free medium for 1 h. The collected medium was cleared by centrifugation at 10’000 rcf for 3 min at 4°C. Proteins were precipitated by the addition of 0.25 volumes trichloroacetic acid (TCA), pelleted at 20’000 rcf at 4°C for 15 min and washed with ice-cold acetone twice.

For mass spectrometry, 50 μl 2 M guanidium hydrochloride, 0.2 M HEPES, pH 8.3, 10 mM TCEP was added and the pellet was sonicated in a Bioruptor Pico cooled by Minichiller 300 (both Diagenode). Chloroacetamide was added to a final concentration of 15 mM and samples were incubated for 10 min at 95°C with gentle agitation. The concentration of guanidium hydrochloride was reduced to 0.5 M by dilution with 100 mM ammonium bicarbonate and the samples were digested with 0.5 μg sequencing grade trypsin (Promega) over-night at 37 °C. The peptides were purified using C18 columns (BioPureSPN Mini Proto 300 C18, Nest Group) according to the manufacturer’s instructions and dried. Detailed information for data acquisition by mass spectrometry and the analysis can be found in supplemental experimental procedures. Gene Ontology (GOterm) enrichment analysis was performed using GORILLA (Eden et al, 2009; Eden et al, 2007).

For SDS-gel electrophoresis, TCA pellets of secreted proteins and post-nuclear supernatants of cell lysates were boiled in SDS-gel sample buffer at 95°C for 10 min. Gels were stained with Commassie-R or subjected to immunoblot analysis.

### Lentiviral transduction

RNA was isolated from HeLaα cells using the RNeasy Mini Kit (Qiagen) and the RNase-Free DNase Set (Qiagen) and cDNA was reverse-transcribed using the IScript cDNA Synthesis Kit (BioRad). The coding sequences of Arfs were amplified by PCR (primers listed in Suppl. Table S1) and inserted into the pQXCIP plasmid (Takara Bio) using the AgeI and BamHI restriction sites. Plasmids were transfected into the packaging cell line Phoenix-ampho (Nolan lab, Stanford University) using jetPRIME (Polyplus Transfection). After 24 h, medium was exchanged to a smaller volume of medium supplemented with 1 mM pyruvate. Medium containing the viral particles was collected after 36 h and cleared by filtration through a 0.45-μm filter. After addition of polybrene to 20 μg/ml, the supernatant was transferred onto target cells. Selection was started 48 h after transduction using medium containing 1.5 mg/ml puromycin (InvivoGen). After 10 days, single cells were sorted by FACS (FACS AriaIII) into 96-well plates with puromycin-containing growth medium supplemented with 10% conditioned medium. After approximately 14 days, clonal cell lines were expanded and examined.

### Statistics

SuperPlots were generated according to Lord et al (2020) and statistical analysis was done with Prism8 (GraphPad) using unpaired one-way ANOVA and unpaired, two-tailed t-test, respectively.

## Acknowledgments

We thank the Imaging Core Facility (IMCF, University of Basel), in particular Dr. Kai Schleicher, Dr. Alexia Ferrand, and Laurent Guerard, for assistance at the microscopes and with data analysis, and Janine Bögli and Stella Stefanova of the FACS Core Facility for their support. This work was supported by Grant 31003A-182519 from the Swiss National Science Foundation.

## Author contributions

MS and MP designed the experiments. MP performed the majority of the experiments. KB performed the mass spectrometry analysis. CPB performed electron microscopy experiments. MS and MP analyzed the data and wrote the manuscript

## Supplemental Experimental Procedures

### Data acquisition and analysis for Mass Spectrometry

Desalted peptides were resuspended in 0.1% aqueous formic acid and subjected to LC–MS/MS analysis using a Q Exactive Plus Mass Spectrometer coupled with an EASY-nLC 1000 (both Thermo Fisher Scientific) and a custom-made column heater set to 60°C. Peptides were resolved using a RP-HPLC column (75μm × 30cm) packed in-house with C18 resin (ReproSil-Pur C18–AQ, 1.9 μm resin; Dr. Maisch GmbH) at a flow rate of 0.2 μl/min. The following stepwise gradient was used for peptide separation: from 5% B to 10% B over 5 min, from 10% B to 35% B over 45 min, from 35% B to 50% B over 10 min, and finally from 50% B to 95% B over 2 min followed by 18 min at 95% B. Buffer A was 0.1% formic acid in water and buffer B was 80% acetonitrile, 0.1% formic acid in water.

The mass spectrometer was operated in DDA mode with a total cycle time of approximately 1 s. Each MS1 scan was followed by high-collision-dissociation (HCD) of the 10 most abundant precursor ions with dynamic exclusion set to 45 seconds. For MS1, 3e6 ions were accumulated in the Orbitrap over a maximum time of 100 ms and scanned at a resolution of 70,000 FWHM (at 200 m/z). MS2 scans were acquired at a target setting of 1e5 ions, maximum accumulation time of 100 ms and a resolution of 35,000 FWHM (at 200 m/z). Singly charged ions and ions with unassigned charge state were excluded from triggering MS2 events. The normalized collision energy was set to 27%, the mass isolation window was set to 1.4 m/z and one microscan was acquired for each spectrum.

The acquired raw-files were imported into the Progenesis QI software (v2.0, Nonlinear Dynamics Limited), which was used to extract peptide precursor ion intensities across all samples applying the default parameters. The generated mgf-file was searched using MASCOT against a human database (consisting of 41484 forward and reverse protein sequences downloaded from Uniprot on 20200417) and 392 commonly observed contaminants using the following search criteria: full tryptic specificity was required (cleavage after lysine or arginine residues, unless followed by proline); 3 missed cleavages were allowed; carbamidomethylation (C) was set as fixed modification; oxidation (M) and acetyl (Protein N-term) were applied as variable modifications; mass tolerance of 10 ppm (precursor) and 0.02 Da (fragments). The database search results were filtered using the ion score to set the false discovery rate (FDR) to 1% on the peptide and protein level, respectively, based on the number of reverse protein sequence hits in the dataset. Quantitative analysis results from label-free quantification were processed using the SafeQuant R package v.2.3.2. (PMID:27345528) to obtain peptide relative abundances. This analysis included global data normalization by equalizing the total peak/reporter areas across all LC-MS runs, data imputation using the knn algorithm, summation of peak areas per protein and LC-MS/MS run, followed by calculation of peptide abundance ratios. Only isoform specific peptide ion signals were considered for quantification. To meet additional assumptions (normality and homoscedasticity) underlying the use of linear regression models and t-Tests, MS-intensity signals were transformed from the linear to the log-scale. The summarized peptide expression values were used for statistical testing of between condition differentially abundant peptides. Here, empirical Bayes moderated t-Tests were applied, as implemented in the R/Bioconductor limma package(PMID: 25605792). The resulting per protein and condition comparison p-values were adjusted for multiple testing using the Benjamini-Hochberg method.

**Suppl. Figure S1:**
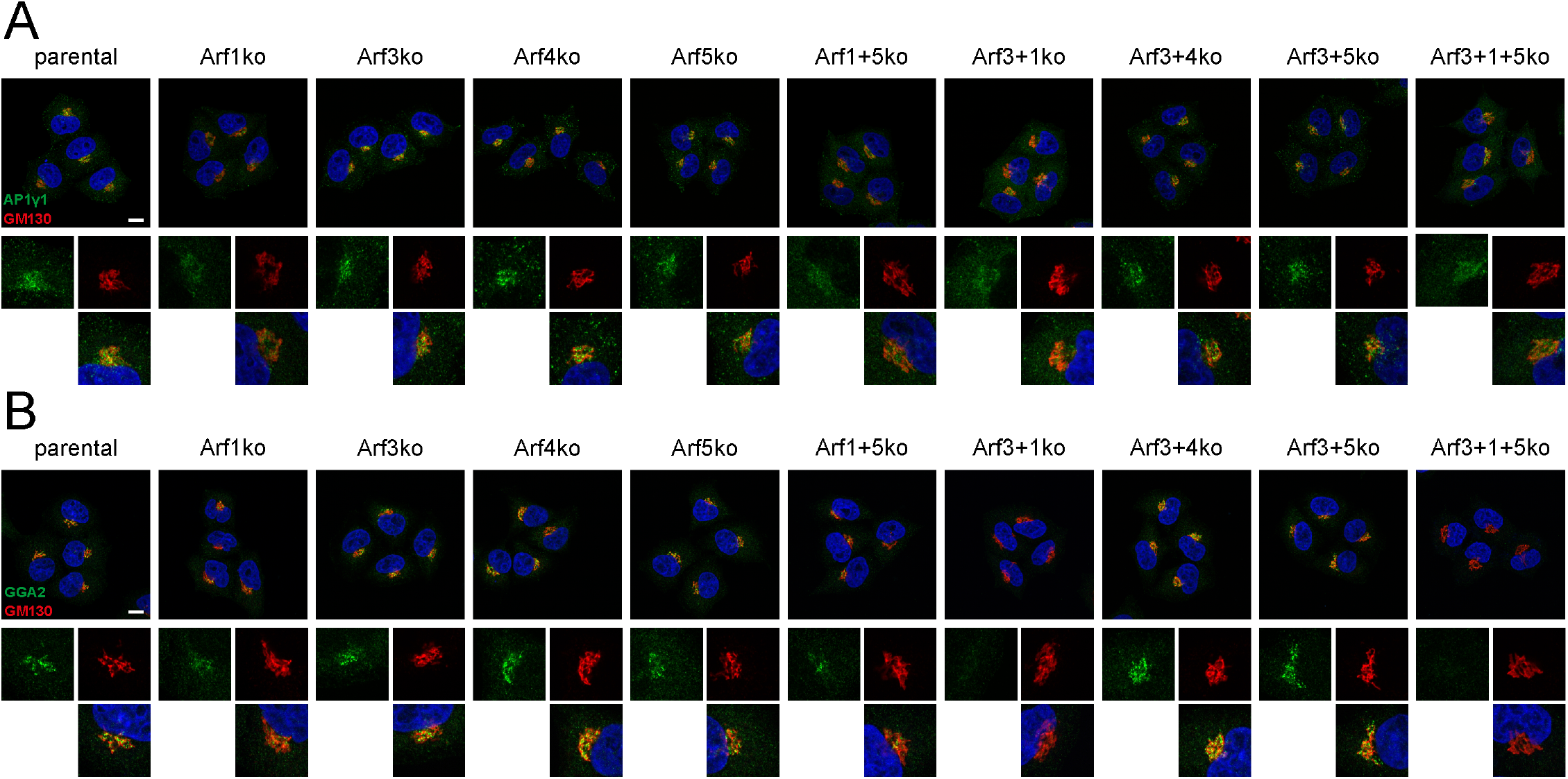
Immunostaining for the Golgi marker GM130 and coat proteins. Parental HeLaα cells and all Arf knockout cell lines were co-immunostained for GM130 (red) and AP1γ1 (A) or GGA2 (B) (green), respectively. DAPI signal shown in blue. Lower panels show magnified image sections (5x). Scale bar, 10 μm.

**Supplementary Table S1:**
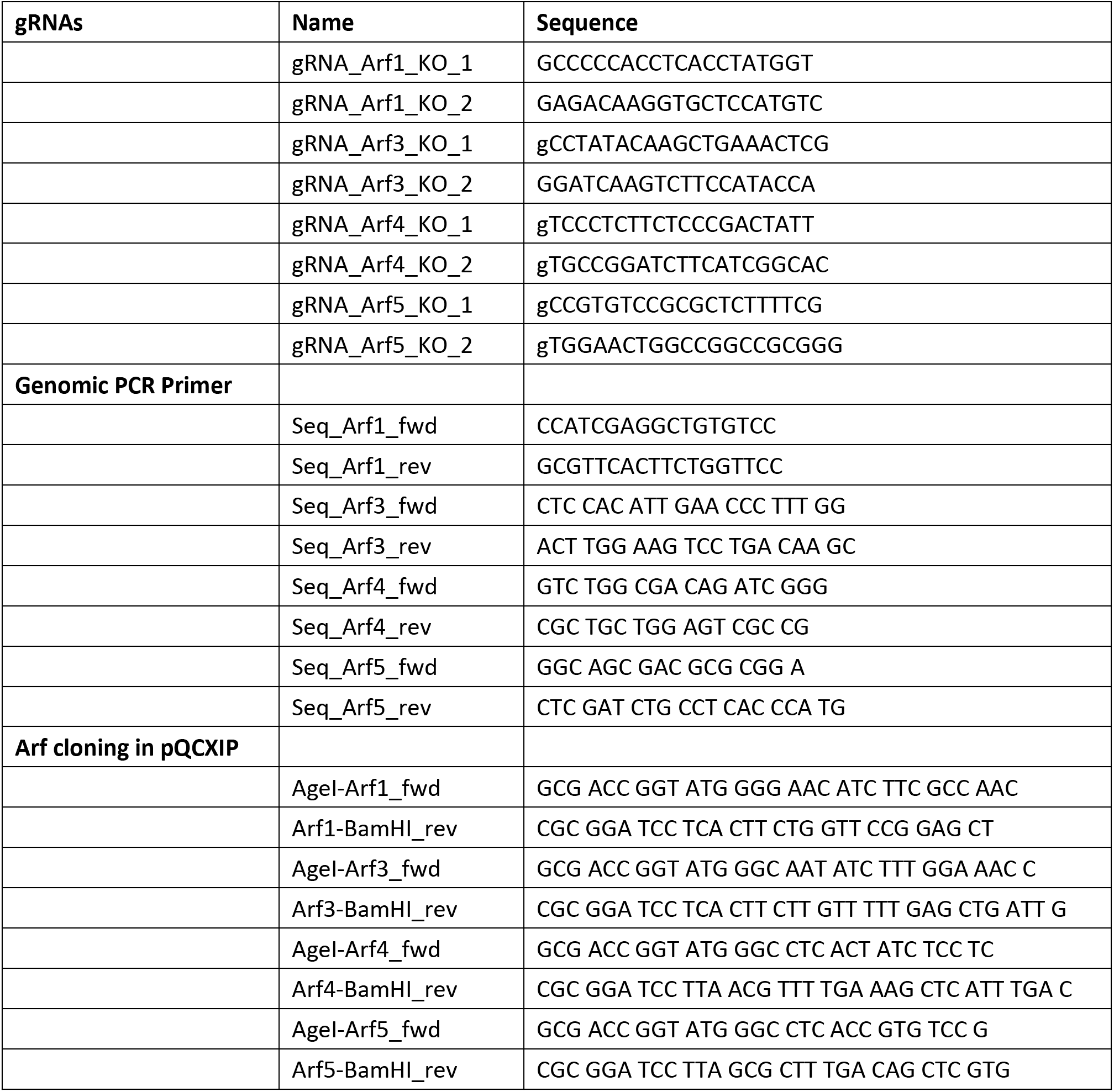
Sequences of gRNAs and PCR primers.

**Supplementary Table S2:**
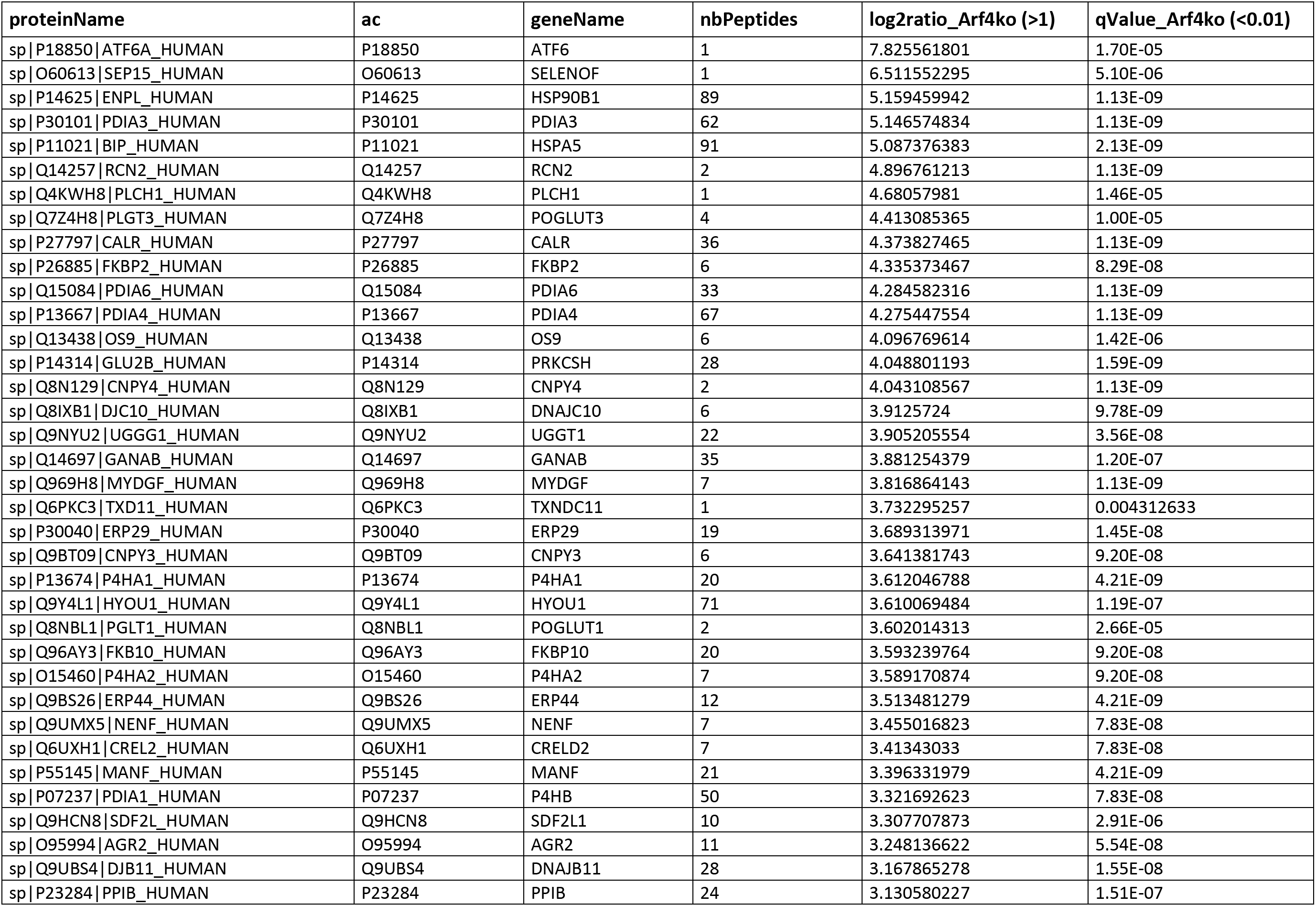
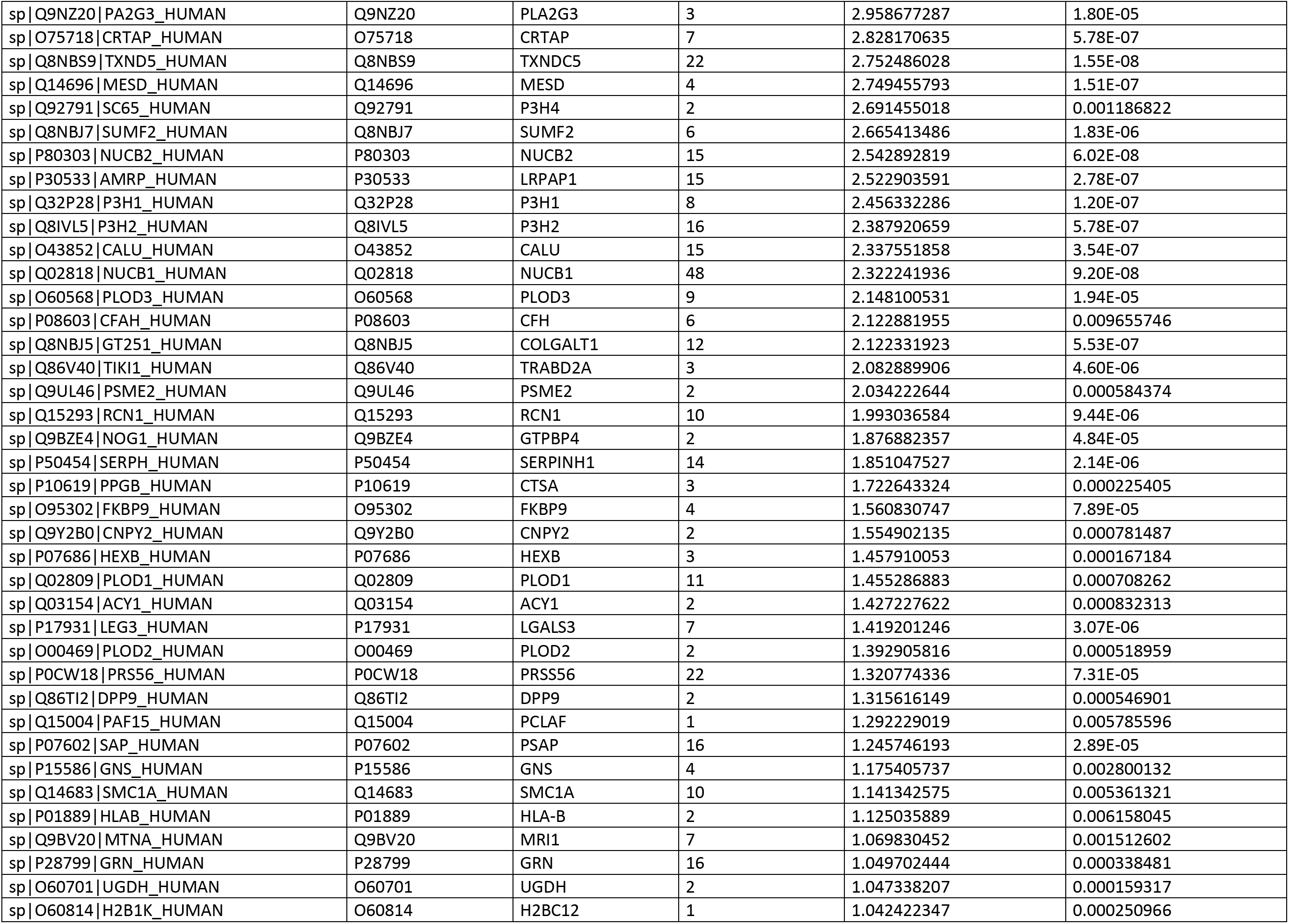
Significant up-regulated hits in the secretome of Arf4ko vs parental HeLaα cells.

**Supplementary Table S3:**
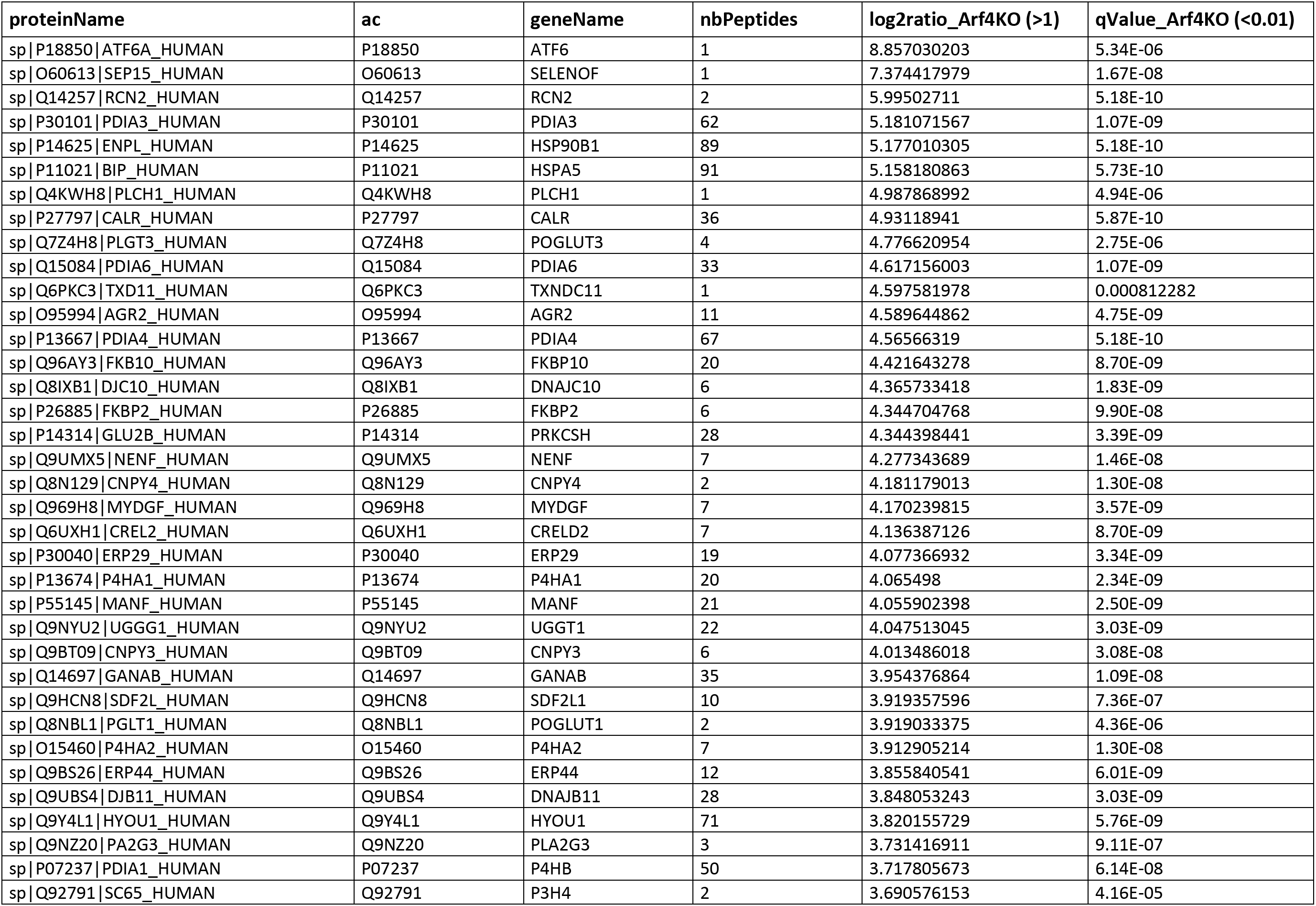
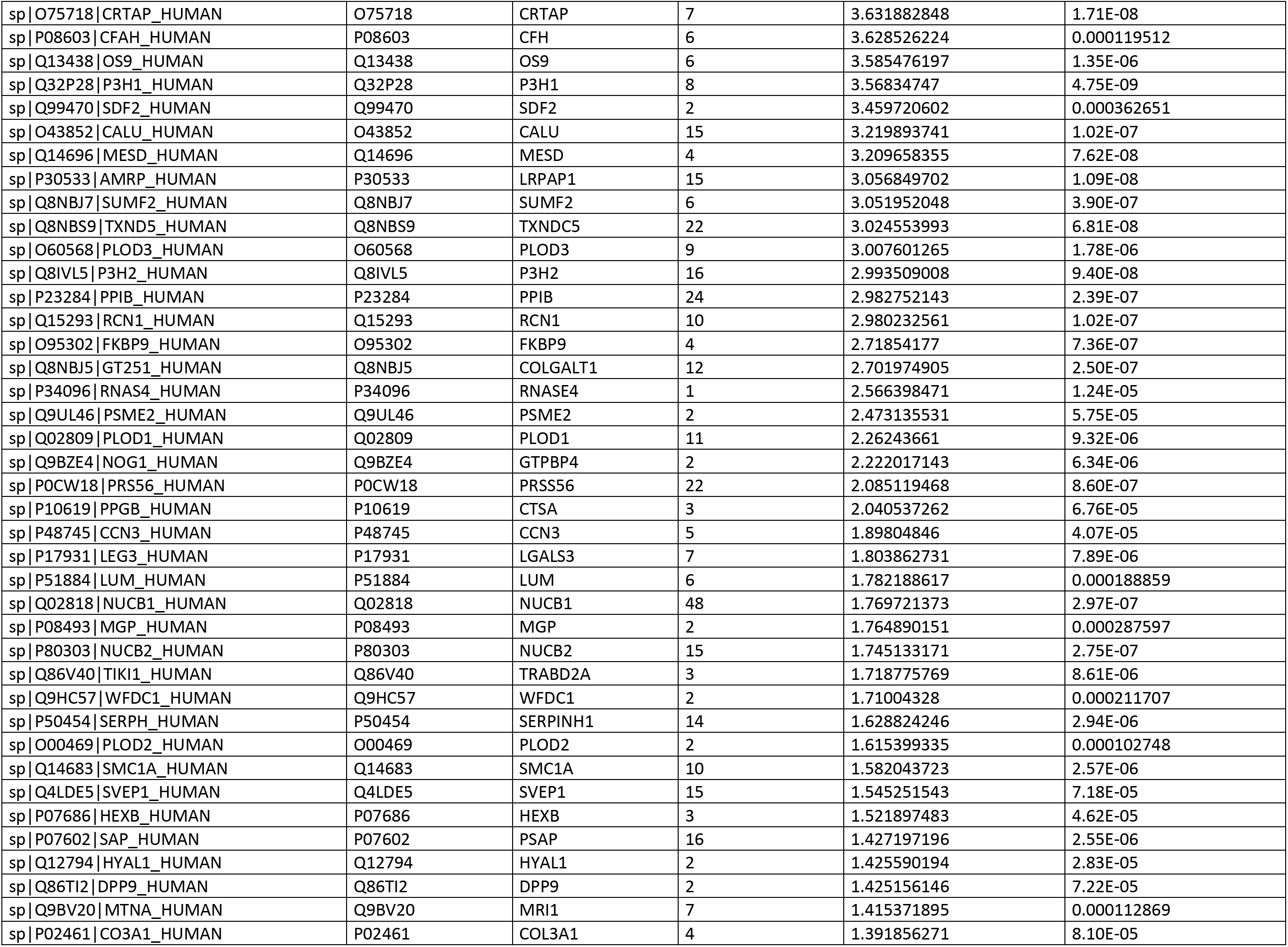
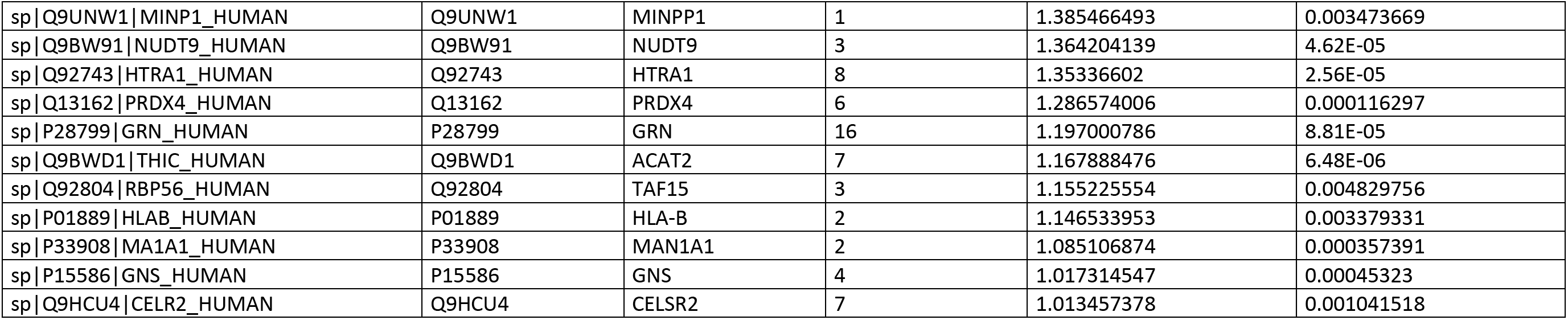
Significant up-regulated hits in the secretome of Arf3+4ko vs parental HeLaα cells.

**Supplementary Table S4:**
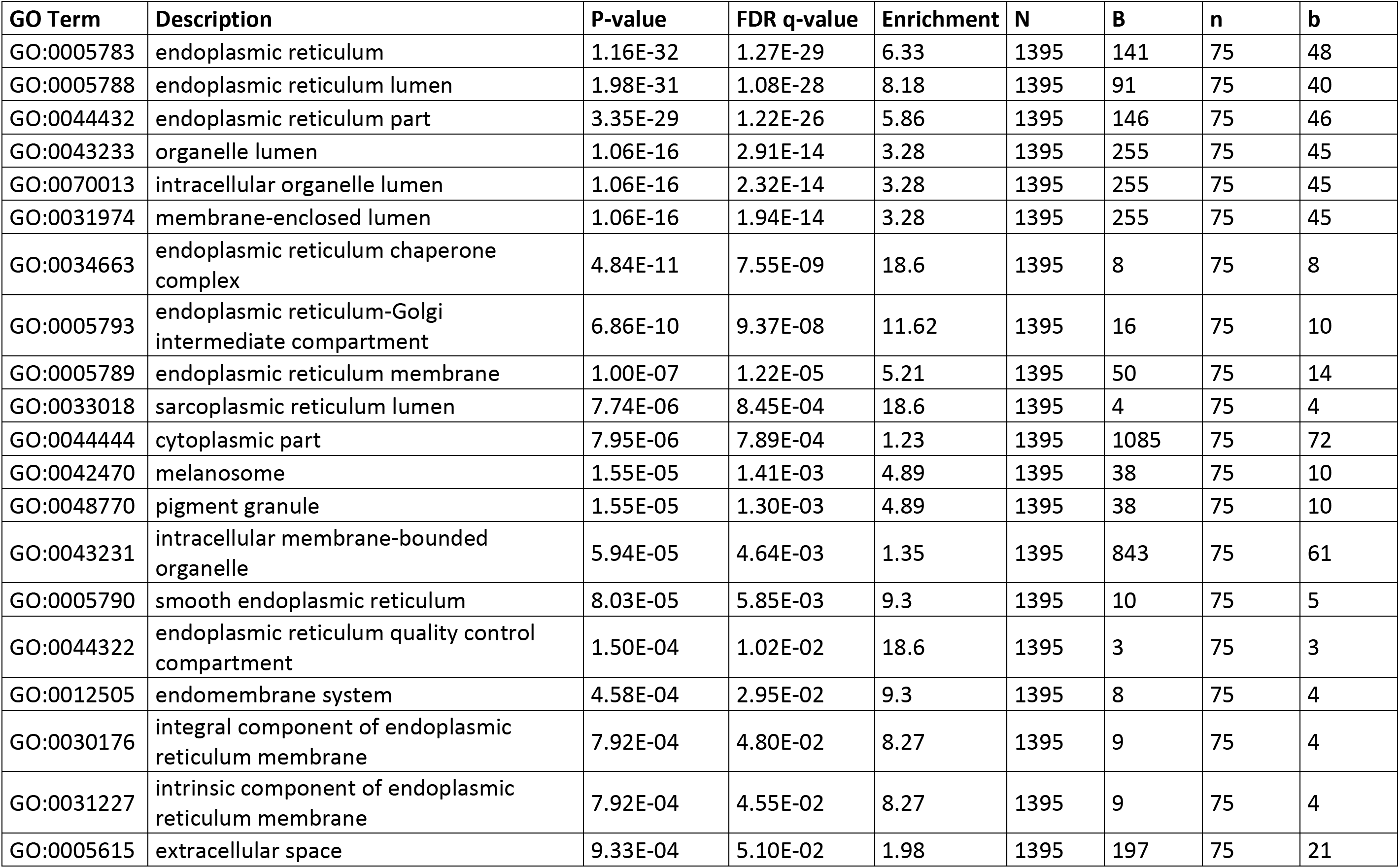
Enriched GOterms for secretome of Arf4ko vs parental HeLaα cells. N is the total number of genes; B is the total number of genes associated with a specific GO term; n is the flexible cutoff;

| GO Term | Description | P-value | FDR q-value | Enrichment | N | B | n | b |
| --- | --- | --- | --- | --- | --- | --- | --- | --- |
| GO:0005783 | endoplasmic reticulum | 1.16E-32 | 1.27E-29 | 6.33 | 1395 | 141 | 75 | 48 |
| GO:0005788 | endoplasmic reticulum lumen | 1.98E-31 | 1.08E-28 | 8.18 | 1395 | 91 | 75 | 40 |
| GO:0044432 | endoplasmic reticulum part | 3.35E-29 | 1.22E-26 | 5.86 | 1395 | 146 | 75 | 46 |
| GO:0043233 | organelle lumen | 1.06E-16 | 2.91E-14 | 3.28 | 1395 | 255 | 75 | 45 |
| GO:0070013 | intracellular organelle lumen | 1.06E-16 | 2.32E-14 | 3.28 | 1395 | 255 | 75 | 45 |
| GO:0031974 | membrane-enclosed lumen | 1.06E-16 | 1.94E-14 | 3.28 | 1395 | 255 | 75 | 45 |
| GO:0034663 | endoplasmic reticulum chaperone complex | 4.84E-11 | 7.55E-09 | 18.6 | 1395 | 8 | 75 | 8 |
| GO:0005793 | endoplasmic reticulum-Golgi intermediate compartment | 6.86E-10 | 9.37E-08 | 11.62 | 1395 | 16 | 75 | 10 |
| GO:0005789 | endoplasmic reticulum membrane | 1.00E-07 | 1.22E-05 | 5.21 | 1395 | 50 | 75 | 14 |
| GO:0033018 | sarcoplasmic reticulum lumen | 7.74E-06 | 8.45E-04 | 18.6 | 1395 | 4 | 75 | 4 |
| GO:0044444 | cytoplasmic part | 7.95E-06 | 7.89E-04 | 1.23 | 1395 | 1085 | 75 | 72 |
| GO:0042470 | melanosome | 1.55E-05 | 1.41E-03 | 4.89 | 1395 | 38 | 75 | 10 |
| GO:0048770 | pigment granule | 1.55E-05 | 1.30E-03 | 4.89 | 1395 | 38 | 75 | 10 |
| GO:0043231 | intracellular membrane-bounded organelle | 5.94E-05 | 4.64E-03 | 1.35 | 1395 | 843 | 75 | 61 |
| GO:0005790 | smooth endoplasmic reticulum | 8.03E-05 | 5.85E-03 | 9.3 | 1395 | 10 | 75 | 5 |
| GO:0044322 | endoplasmic reticulum quality control compartment | 1.50E-04 | 1.02E-02 | 18.6 | 1395 | 3 | 75 | 3 |
| GO:0012505 | endomembrane system | 4.58E-04 | 2.95E-02 | 9.3 | 1395 | 8 | 75 | 4 |
| GO:0030176 | integral component of endoplasmic reticulum membrane | 7.92E-04 | 4.80E-02 | 8.27 | 1395 | 9 | 75 | 4 |
| GO:0031227 | intrinsic component of endoplasmic reticulum membrane | 7.92E-04 | 4.55E-02 | 8.27 | 1395 | 9 | 75 | 4 |
| GO:0005615 | extracellular space | 9.33E-04 | 5.10E-02 | 1.98 | 1395 | 197 | 75 | 21 |

**Supplementary Table S5:** Enriched GOterms for secretome of Arf3+4ko vs parental HeLaα cells. N is the total number of genes; B is the total number of genes associated with a specific GO term; n is the flexible cutoff;

| GO Term | Description | P-value | FDR q-value | Enrichment | N | B | n | b |
| --- | --- | --- | --- | --- | --- | --- | --- | --- |
| GO:0005783 | endoplasmic reticulum | 7.75E-31 | 8.46E-28 | 5.69 | 1395 | 141 | 87 | 50 |
| GO:0005788 | endoplasmic reticulum lumen | 8.49E-31 | 4.63E-28 | 7.4 | 1395 | 91 | 87 | 42 |
| GO:0044432 | endoplasmic reticulum part | 1.45E-27 | 5.28E-25 | 5.27 | 1395 | 146 | 87 | 48 |
| GO:0043233 | organelle lumen | 2.77E-18 | 7.55E-16 | 3.21 | 1395 | 255 | 87 | 51 |
| GO:0070013 | intracellular organelle lumen | 2.77E-18 | 6.04E-16 | 3.21 | 1395 | 255 | 87 | 51 |
| GO:0031974 | membrane-enclosed lumen | 2.77E-18 | 5.03E-16 | 3.21 | 1395 | 255 | 87 | 51 |
| GO:0005793 | endoplasmic reticulum-Golgi intermediate compartment | 1.01E-10 | 1.57E-08 | 11.02 | 1395 | 16 | 87 | 11 |
| GO:0034663 | endoplasmic reticulum chaperone complex | 1.68E-10 | 2.29E-08 | 16.03 | 1395 | 8 | 87 | 8 |
| GO:0005789 | endoplasmic reticulum membrane | 7.13E-07 | 8.65E-05 | 4.49 | 1395 | 50 | 87 | 14 |
| GO:0005615 | extracellular space | 6.75E-06 | 7.37E-04 | 2.28 | 1395 | 197 | 87 | 28 |
| GO:0005790 | smooth endoplasmic reticulum | 8.53E-06 | 8.47E-04 | 9.62 | 1395 | 10 | 87 | 6 |
| GO:0033018 | sarcoplasmic reticulum lumen | 1.42E-05 | 1.29E-03 | 16.03 | 1395 | 4 | 87 | 4 |
| GO:0042470 | melanosome | 5.92E-05 | 4.98E-03 | 4.22 | 1395 | 38 | 87 | 10 |
| GO:0048770 | pigment granule | 5.92E-05 | 4.62E-03 | 4.22 | 1395 | 38 | 87 | 10 |
| GO:0044322 | endoplasmic reticulum quality control compartment | 2.35E-04 | 1.71E-02 | 16.03 | 1395 | 3 | 87 | 3 |
| GO:0043202 | lysosomal lumen | 7.17E-04 | 4.90E-02 | 4.32 | 1395 | 26 | 87 | 7 |
| GO:0012505 | endomembrane system | 8.16E-04 | 5.24E-02 | 8.02 | 1395 | 8 | 87 | 4 |

